# Human HYPOMAP: A comprehensive spatio-cellular map of the human hypothalamus

**DOI:** 10.1101/2023.09.15.557967

**Authors:** John A. Tadross, Lukas Steuernagel, Georgina K.C. Dowsett, Katherine A. Kentistou, Sofia Lundh, Marta Porniece-Kumar, Paul Klemm, Kara Rainbow, Henning Hvid, Katarzyna Kania, Joseph Polex-Wolf, Lotte Bjerre-Knudsen, Charles Pyke, John R. B. Perry, Brian Y.H. Lam, Jens C. Brüning, Giles S.H. Yeo

## Abstract

The hypothalamus is a brain region that plays a key role in coordinating fundamental biological functions. However, our understanding of the underlying cellular components and circuitry, have, until recently, emerged primarily from rodent studies. Here, we combine a single-nucleus sequencing database of 433,369 human hypothalamic cells, with spatial transcriptomics, to present a comprehensive spatio-cellular transcriptional map of the human hypothalamus, the ‘HYPOMAP’. Analysing hypothalamic leptin melanocortin pathway neuronal populations that play a role in appetite control, we identify spatially distinct populations of arcuate nucleus *POMC* and *AGRP* neurons, and their receptors *MC3R* and *MC4R*. Next, we map the cells expressing incretin receptors, targets of the new generation of anti-obesity medications, and uncover transcriptionally distinct *GLP1R* and *GIPR*-expressing cellular populations. Finally, out of the 458 hypothalamic cell types in HYPOMAP, we find 182 neuronal clusters are significantly enriched in expression of BMI GWAS genes. This enrichment is driven by 375 ‘effector’ genes, with rare deleterious variants in 6 of these; *MC4R*, *PCSK1*, *POMC*, *CALCR*, *BSN* and *CORO1A*, the last of which has previously not been linked to obesity; being significantly associated with changes in BMI at the population level. Thus, the HYPOMAP provides a detailed atlas of the human hypothalamus in a spatial context, and serves as an important resource to identify novel druggable targets for treating a wide range of conditions, including reproductive, circadian, and metabolic disorders.

## Introduction

The hypothalamus is a brain region that plays a key role in coordinating fundamental biological functions, including maintaining body temperature, sleep, thirst, and energy homeostasis, as well as regulating sexual and parental behaviour, response to stress, and circadian rhythms. Yet in spite of its importance, the inaccessibility of the human hypothalamus has meant that, to date, our understanding of its architecture, has emerged primarily from rodent studies.

For instance, while human genetic studies have uncovered many of the major components of the hypothalamic appetitive ^1^ and reproductive ^2^ control pathways, it has only been possible to illuminate their underlying cellular components and circuitry through detailed neuroanatomical mapping studies in mice. The fat sensing leptin-melanocortin pathway, which comprises agouti-related peptide (AgRP)- and pro-opiomelanocortin (POMC)-expressing neurons in the hypothalamic arcuate nucleus (ARC), acting through intra- and extrahypothalamic projections to control food intake and energy expenditure, is a case in point. We know that it plays a key role in the control of appetite because genetic disruption at all levels of the pathway results in severe obesity, not only in humans and mice, but in many other vertebrate species as well ^1,3^. Recently we found that the leptin-melanocortin pathway also plays important roles in linear growth and pubertal onset, *via* the melanocortin 3 receptor (MC3R) ^4^. However, our understanding of the melanocortin neurocircuitry is largely derived from functional studies in mice ^1,3^.

Despite the paucity of knowledge about the human hypothalamic circuitry, currently licensed therapies that reduce food intake, including semaglutide ^5^ and tirzepatide ^6^, primarily target the hypothalamus ^7^ and hindbrain ^8^. Semaglutide is a glucagon-like peptide-1 receptor (GLP1R) agonist, and tirzepatide a GLP1R/gastric inhibitory polypeptide receptor (GIPR) co-agonist, and both are thought to mediate their effects on appetite, at least in part, *via* POMC neurons ^7^. However, studies supporting these modes of action are largely derived from studies in mice ^7,9^. Additionally, setmelanotide, an MC4R agonist that targets the melanocortin system itself, has recently been approved for rare genetic causes of obesity ^10^. Given the therapeutic focus on the hypothalamus, enhancing our understanding of its human-specific architecture is crucial.

Over the past decade, access to fresh frozen and fixed human donor brain samples from brain banks has markedly improved ^11^. These precious samples, coupled with developments in droplet single-cell/nucleus sequencing and spatial transcriptomic technologies, have provided us the opportunity to map the functional architecture of the human hypothalamus. Following on from our generation of a unified murine hypothalamic single cell database ^12^, here, we have integrated single nucleus RNA sequencing (snRNAseq) data from 433,369 hypothalamic cells with spatial transcriptomics to create a comprehensive spatio-cellular map of the human hypothalamus, the HYPOMAP. This resource has allowed us to uncover previously unappreciated spatial heterogeneity of non-neuronal hypothalamic cell-types, and provided the opportunity to delineate the spatial context of neuronal populations, such as those expressing components of the leptin-melanocortin pathway and GLP1/GIP receptors, both targets of current anti-obesity therapeutics. Finally, HYPOMAP has provided a framework for the integration of population level genetic data, which we show here for BMI associated genes, allowing us to identify new components in the regulation of body-weight. Thus, this new open-access resource will be vital in our understanding of the human hypothalamus and facilitate the identification of novel druggable targets for treating a wide range of conditions, including reproductive, circadian, and metabolic disorders.

## Results

### HYPOMAP captures 433K human hypothalamic cells

We collected fresh frozen hypothalamic tissue from eight brain donors of normal BMI (range 18-28, clinical data shown **Supp Table 1**), and performed snRNASeq (see Methods). We and others have previously shown snRNASeq produces comparable data to single-cell RNA sequencing (scRNASeq) ^12,13^, thus allowing us to work with frozen archival samples. After quality control steps (see Methods), we captured 311,964 nuclei with an average of 4541 ± 10.6 (mean ± SEM) counts detected per nucleus (mean 2040 ± 2.4 genes per nucleus). In addition, we extracted the expression matrix for 121,405 nuclei (20,331 ± 55.9 [mean ± SEM] counts detected and 5097 ± 7.1 genes per nucleus) from the hypothalamic regions of three separate donors, captured in the publicly available whole-brain dataset (Siletti et al, https://doi.org/10.1101/2022.10.12.511898) (**Supp Table 1**). We integrated the two datasets *via* scvi-tools to correct for donor and study heterogeneity ^12,14^, and generated a reference single-cell database of the human hypothalamus consisting of a total of 433,369 nuclei, we call the human ‘HYPOMAP’ (**Extended Data Figure 1, 2**). A uniform manifold approximation plot (UMAP) is shown in **Figure 1A**, illustrating the different major cell types identified from the human hypothalamus. Here, we examined the expression of key transcription factors, used as regional markers, demonstrating our dataset indeed spans the hypothalamus (**Extended Data Figure 3**).

**Figure 1:**
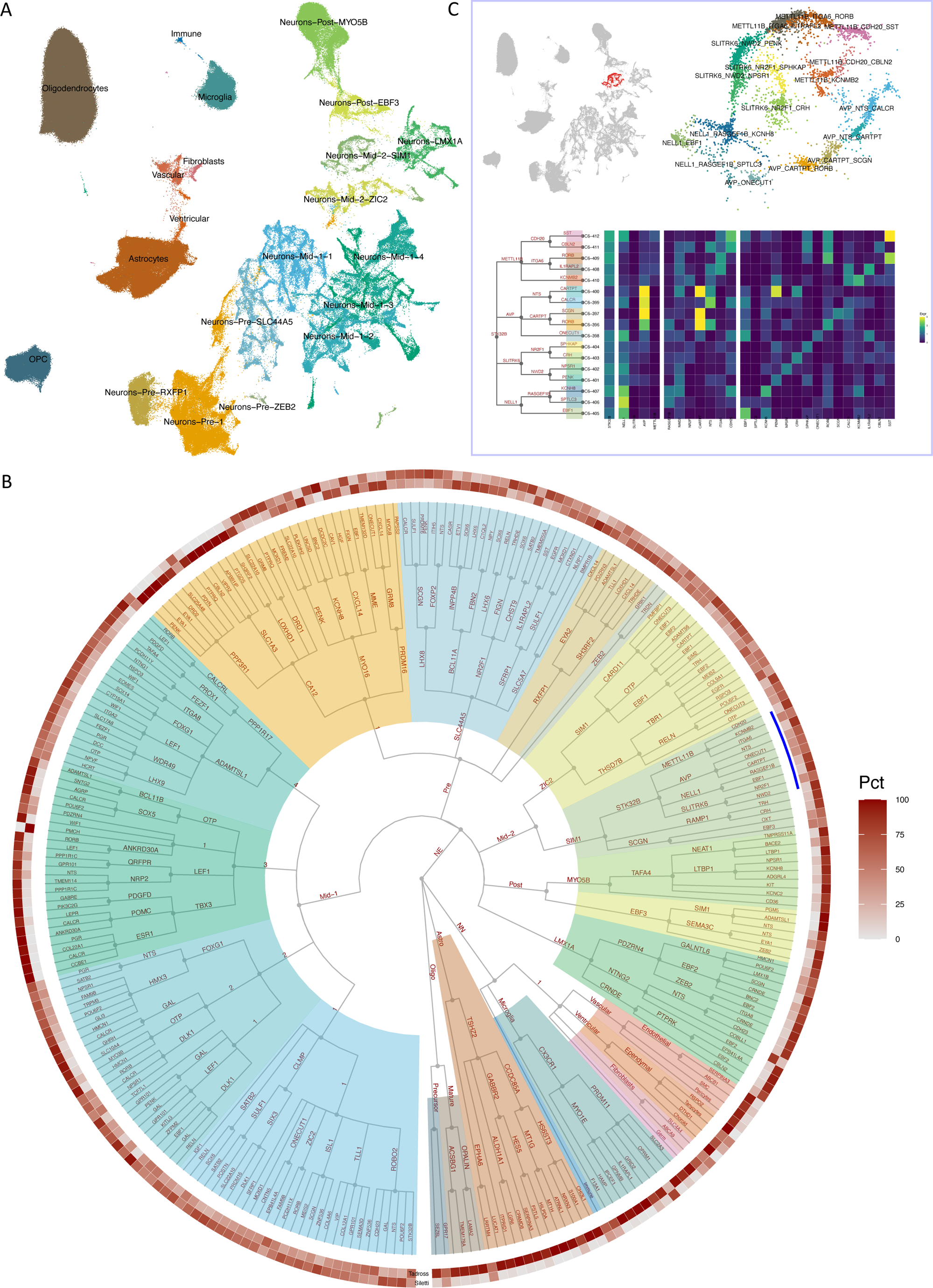
Integrated single-nucleus RNA sequencing (snRNAseq) reference atlas of the human hypothalamus. **a)** Uniform Manifold Approximation and Projection (UMAP) plot of the integrated atlas colored by major cell classes. Coloring corresponds to background colors in (b). scvi-tools was utilized to integrate the heterogeneous human samples originating from different donors and studies. **b)** Unbiased clustering of the integrated human snRNAseq data. For 4 major subgroups, neurons, astrocytes, oligodendrocytes and other non-neuronal cells, multiple flat clustering using the leiden algorithm were generated and combined into a hierarchical consensus tree. The final tree spans 7 different levels (C0-C6) with 4 to 458 distinct clusters, however, to improve visibility the circular tree only includes the first 6 levels with up to 265 clusters. Nodes in the tree were anntotated using the most informative marker genes, which are shown on the edges of the circular tree. The background color marks major cell groups and corresponds to the coloring of clusters in the UMAP plot of (a). The heatmap ring depicts the relative contribution of cells by the two included studies to each cluster on level C5 **c)** Details on cluster NE_Mid2_SIM1_STK32B (C3) neurons highlighted in blue in the circular tree in (b). This cluster includes the magnocellular AVP neurons as well as multiple SST-expressing subclusters. They are unified by their expression of the kinase STK32B. The UMAP plots on the left show the position on the global UMAP and a subset-UMAP containing only cells of the cluster colored by level C6. The coloring of clusters corresponds to the dendrogram shown on the right, which gives a detailed overview of the subcluster structure. See the description of (b) for more details on the tree generation and annotation. The heatmap shows the average expression of marker genes used for annotation (edges of the dendrogram).

### Flexible cell-type classification via multi-level hierarchical clustering

HYPOMAP comprises 166,475 neurons (NE), 175,109 oligodendrocytes (Oligo), 61,579 astrocytes (Astro) and 30,206 cells from other non-neuronal (NN) cell types (**Figure 1A**). Here, we adopted a multi-level (granularity) hierarchical clustering strategy by coupling leiden and multi-resolution tree (mrtree) algorithms, to enable flexibility in cell-type classification and ensure optimal granularity for downstream analyses of specific populations (see Methods for details). The final circular dendrogram (tree) consists of 7 levels and 458 clusters (levels C0-C5 in **Figure 1B** to preserve visibility, C0=4, C1=12, C2=29, C3=59, C4=134, C5=265, C6=458). Cluster nomenclature was determined based on the most informative marker gene exhibiting the largest global differences to all clusters, and local differences to its sibling clusters; if a marker gene could not be determined, clusters were manually named with their common names, or a sequential number. All marker genes are labelled on the tree edges (**Figure 1B**). At cluster level C1, hypothalamic neurons are segregated into 5 sub-populations, largely following their broad anatomical origin, from the preoptic/anterior (Pre), middle (Mid) to the posterior (Post) hypothalamus (**Figure 1A**). Mid hypothalamic neurons exhibit a higher level of heterogeneity compared to other regions, and a larger number of clusters in the tree (**Figure 1A,B**).

To illustrate the functionality of our atlas, we have focused on a subset of *SIM1* neurons at level C3 as an exemplar, highlighted in blue in **Figure 1B** and red in **Figure 1C** (inset). These neuronal populations are marked with expression of *STK32B* (**Figure 1C**). They are segregated into 4 different classes at the next level C4, one of which has high expression of *AVP*. Moving to level C6, and these neurons are divided into 5 further populations, driven by the expression of *NTS*, *CARTPT*, *CALCR*, *SCGN*, *RORB* and *ONECUT*, all showing distinct clustering on the UMAP (**Figure 1C**).

### Spatial cellular mapping via the integration of spatial transcriptomics and snRNAseq

Next, using Visium technology (10X Genomics), we performed high-resolution spatial transcriptomic profiling of 7 hypothalamic sections from 5 donors, covering the preoptic/anterior, middle and the posterior hypothalamus (regional annotation shown in **Figure 2A**). An example of the data-output can be seen in a mid-hypothalamic section (the third section in **Figure 2A**), where we can discriminate spatially-restricted expression of VGLUT2 (*SLC17A6*) and VGAT (*SLC32A1*), which corresponds to the excitatory glutamatergic neurons concentrated within the ventromedial hypothalamus (VMH), and the inhibitory GABA-ergic neurons in the arcuate nucleus (ARC) respectively (**Figure 2B**). The region-specific expression of transcription factors *TBX3*, *FEZF1*, and *SIM1*, which mark the ARC, VMH and PVN respectively on the same section, are also clearly illustrated (**Figure 2C**).

**Figure 2:**
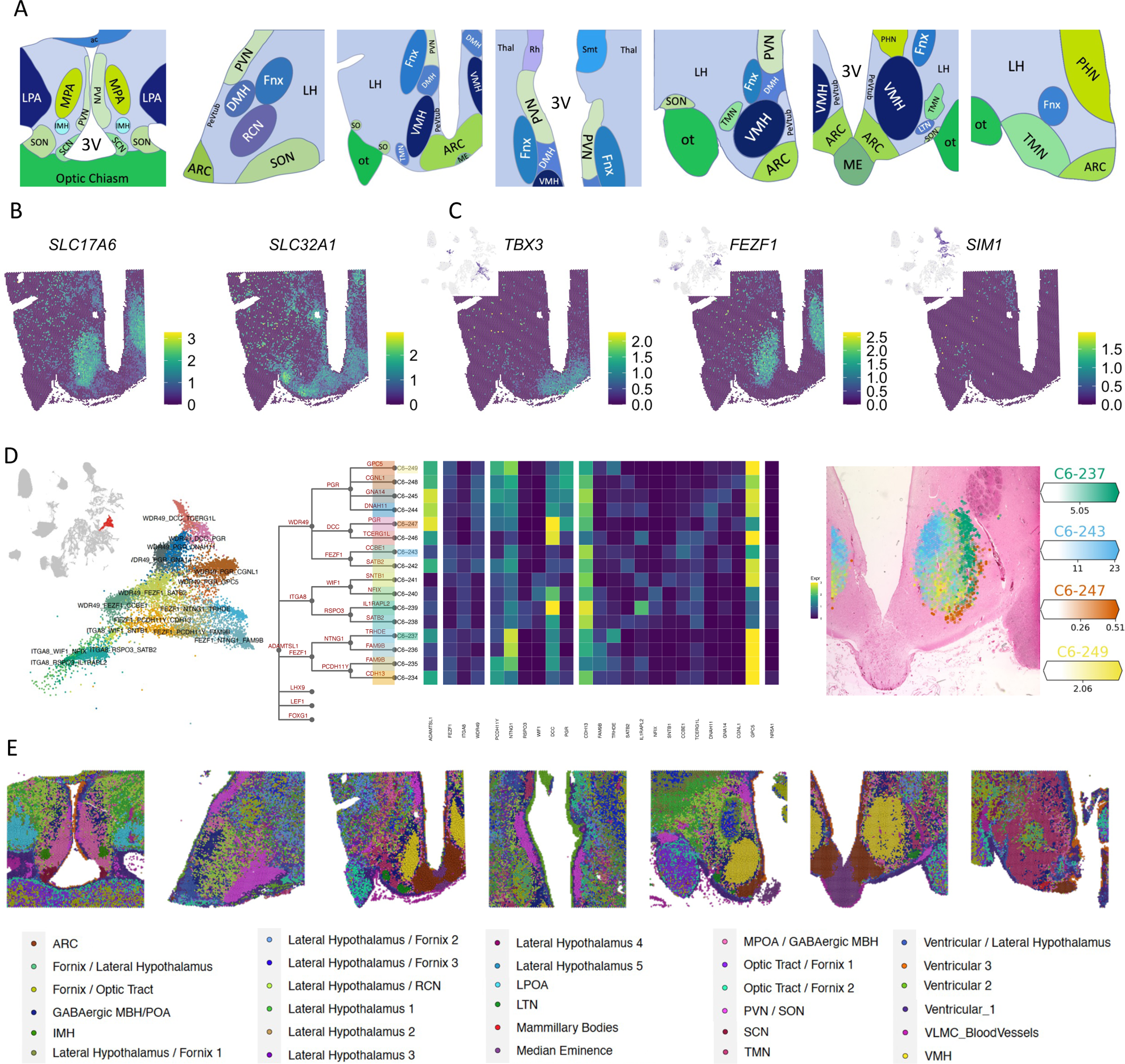
Spatial Transcriptomics of the human hypothalamus and mapping of cell clusters identified by snRNAseq. **a)** A reference atlas diagram of the 7 human hypothalamic sections used in spatial transcriptomics experiment (5 donors). Diagrams were created using the H&E tissue sections, expression of canonical marker genes for hypothalamic nuclei, and a human brain reference atlas in combination. Sections are ordered from most anterior (left) to most posterior (right). LPA = lateral preoptic area; MPA = medial preoptic area; PVN = paraventricular nucleus; IMH = intermediate hypothalamic nucleus; SCN = suprachiasmatic nucleus; SON/SO = supraoptic nucleus; ac = anterior commissure; 3V = third ventricle; ARC = arcuate nucleus; DMH = dorsomedial hypothalamus; RCN = retrochiasmatic nucleus; PeVtub = Periventricular tuberal nucleus; Fnx = fornix; LH = lateral hypothalamus; ot = optic tract; TMN = tuberomammillary nucleus; VMH = ventromedial hypothalamus; ME = median eminence; Thal = thalamus; Rh = rhomboid nucleus of the thalamus; Smt = stria medullaris of thalamus; PHN = posterior hypothalamic nucleus. **b)** Log-normalized expression of SLC17A6, and SLC32A1 (glutamatergic and GABAergic markers respectively) in a medial human hypothalamic section. SLC32A1 shows clear expression in the arcuate nucleus, and the periventricular nucleus. SLC17A6 is expressed in the ventromedial hypothalamus. **c)** Spatial expression plots of transcription factors TBX3, FEZF1 and SIM1, used to mark the ARC, VMH and PVN respectively. Expression of these genes in the snRNAseq dataset also represented. **d)** Details on 3 branches of C3-18 NE_Mid-1_4_ADAMTSL1 (C4-51: NE_Mid-1_4_ADAMTSL1_WDR49, C4-50: NE_Mid-1_4_ADAMTSL1_ITGA8, C4-49: NE_Mid-1_4_ADAMTSL1_FEZF1). (Left) clusters highlighted in the global UMAP and a subset of the UMAP containing only cells of the cluster coloured by C6. The colouring of the clusters corresponds to the dendrogram (middle) which provides an overview of the subcluster structure and displays a heatmap of the average expression of each marker gene in each cluster. (Right) Cell2Location mapping of 4 clusters (highlighted in the dendrogram) at C6 level demonstrates mapping to subregions of the ventromedial hypothalamus in a mid-posterior section of the hypothalamus. **e)** Spatial plots displaying leiden clustering of spots from the co-occurance of snRNAseq cluster abundance scoares. The matrix of cell abundance values of each snRNAseq cluster in each spot was used in the Leiden clustering analysis. Spots in the same cluster will show similar patterns of snRNAseq cell abundance. Clusters were named based on which regions of the hypothalamus they mostly appeared in.

Because Visium technology does not achieve single-cell resolution (each spot typically covering 1-10 cells), we integrated the spatial transcriptomics and snRNAseq data using cell2location, a Bayesian model-based package ^15,16^, to spatially map cell clusters at all levels in the snRNAseq, to the hypothalamic sections shown in **Figure 2A**. **Figure 2D** showcases the spatial mapping of VMH cell clusters identified in the snRNAseq data. At C6 level, we identify 16 VMH neuronal populations. Here, they all show distinct mapping in the VMH (**Extended Data Figure 4**). 4 clusters, C6-237, C6-243, C6-247 and C6-249 (highlighted in **2D dendrogram**) and their mapping in the VMH are shown in the mediobasal hypothalamus section in **Figure 2D**.

To identify tissue regions in which sets of snRNAseq clusters consistently map to, we performed leiden clustering on the cell abundance values of each cluster (at C6 level) in each Visium spot (**Figure 2E**). Through this, we identified 30 clusters, which were annotated based on the hypothalamic regions in which the majority of spots were located. Of note, we identified clusters for the ARC, VMH, PVN/SON, lateral tuberal nucleus (LTN), median eminence (ME), tuberomammillary nucleus (TMN), as well as clusters in predominantly non-neuronal regions e.g fornix and optic tract. Tables displaying the average cell abundance of each snRNAseq cluster (at levels C1:C6) within each spatial cluster can be found in **Supp Tables 9-14**.

### Distinct spatial mapping of non-neuronal populations in the hypothalamus

Non-neuronal cells such as astrocytes, oligodendrocytes, and tanycytes have been historically understudied. While less heterogeneous than their neuronal counterparts, single cell approaches have still revealed considerable diversity ^17^. Here, we characterised the non-neuronal cell populations in the human hypothalamus. According to the snRNAseq data, hypothalamic astrocytes are broadly divided into two major populations, each marked by *EPHA6* and *TSHZ2*. They further split into 7 sub-populations (C3-47: Astro_EPHA6, C3-48: Astro_TSHZ2_ITPRID1, C3-49: Astro_TSHZ2_ALDH1A1, C3-50: Astro_TSHZ2_GABBR2_HES5, C3-51: Astro_TSHZ2_GABBR2_SERPINA3, C3-52: Astro_TSHZ2_CCDC85A_MT1G, C3-53: Astro_TSHZ2_CCDC85A_HS6ST3) at level C3. These all show spatially distinct patterns of expression in the hypothalamus (**Extended Data Figure 5A**). Similarly, we found distinct populations of oligodendrocytes (C2_29 Oligo_OPALIN) and (C2_28 Oligo_ACSBG1) located in the anterior commissure (ac) and the optic chiasm in the anterior hypothalamus respectively, but they are more intermixed in the lateral preoptic area (LPOA, **Extended Data Figure 5B**). Ependymal cells also showed different localization on the hypothalamic sections. In Extended Data Figure 4C, we show that the 3^rd^ ventricle (3V) was lined with *DTHD1*-expressing ependymal cells (C4-94: NN_1_Ventricular_Ependymal_DTHD1), while the tanycytes (C4-95: NN_1_Ventricular_Ependymal_Tanycytes) were mapped to two locations in the MBH: at the base of the 3V and in the median eminence.

Campbell and colleagues ^18^ previously demonstrated discrete subpopulations of tanycytes in the mouse hypothalamus. While our NucSeq data did not capture sufficient tanycytes (∼100 cells) to differentiate between subtypes in humans, we did observe different patterns of expression of *CRYM* and *FRZB* in the median eminence (**Extended Data Figure 5D**). We additionally looked further into expression of tanycyte and ependymal marker genes in the spatial transcriptomics dataset. We see concentrated levels of expression of *DIO2* and *FZD5* below the third ventricle and in the median eminence region, however expression of *STOML3* and *LPAR3* showed distinct expression in spots lining the entirety of the walls of the third ventricle, suggesting that these are ependymal cell markers (**Extended data Figure 5D**). We confirmed these findings using RNAscope.

### Human hypothalamic leptin-melanocortin system

Insights from human and mouse genetics have illuminated the central role of the hypothalamic leptin-melanocortin pathway in the control of mammalian food intake, with genetic disruption resulting in extreme obesity, and more subtle polymorphic variation influencing the population distribution of body-weight ^1,3^. Here, we examined the transcriptomic profiles, as well as the spatial localisation of leptin-responsive melanocortin neurons in the human hypothalamus.

We first focused on neurons expressing *POMC*, the gene encoding the melanocortin peptides. At the highest granularity level C6, we discovered six clusters of *POMC*-expressing neurons (**Figure 3A**, cut off: *POMC*+ clusters in the 99^th^ percentile), and five out of six clusters are mapped to the ARC (C6-278, C6-279, C6-280, C6-289 and C6-283; extended data figure 6A,B,E). Of these, 3 clusters displayed higher average expression of *POMC* (C6-278, C6-279 and C6-280, highlighted on **Figure 3A**). C6-278 (NE-mid-1_3_TBX3_POMC_LEPR) has the highest level of *POMC* expression (**Figure 3B**), and LEPR is one of the key markers for this cluster and is detected in 80% of neurons, with 68.12% of cells in this cluster co-expressing *POMC* and *LEPR*. The spatially mapped *POMC* clusters show distinct patterns of localisation, with the canonical *POMC/LEPR* neurons located adjacent to the median eminence, and the *POMC/CALCR* neurons closest to the 3^rd^ ventricle (**Figure 3C**).

**Figure 3:**
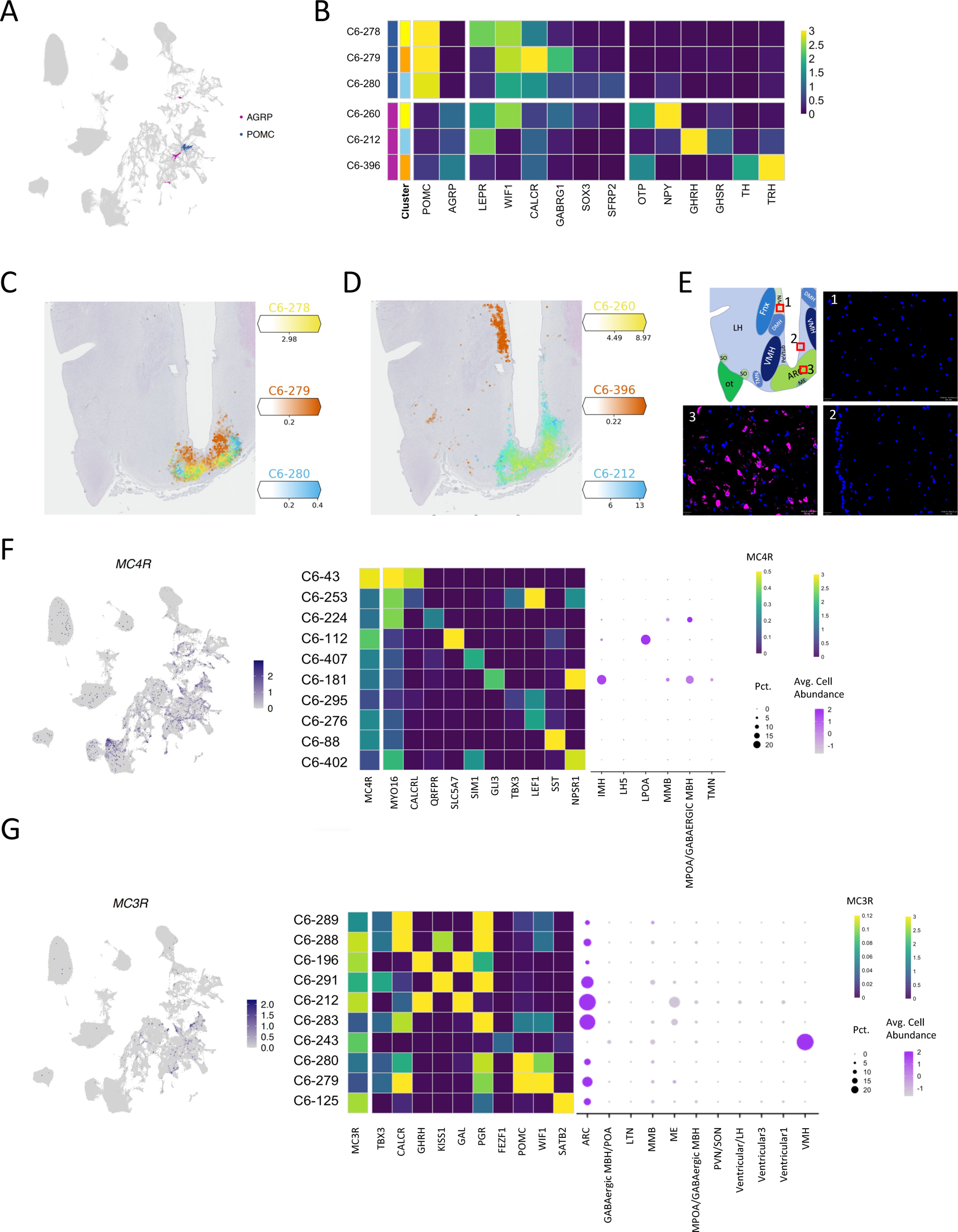
Mapping the leptin-melanocortin pathway. Characterisation of POMC and AGRP neurons in the human hypothalamus. **a)** UMAP plot highlighting 3 clusters with the highest percentage of POMC+ nuclei (blue) and 3 clusters with the highest percentage of AGRP+ nuclei (pink). **b)** Heatmap displaying average log-normalised expression of marker genes for each of the top 3 POMC and AGRP clusters. **c)** Spatial mapping of POMC clusters. Cell abundance scores for C6-278 (yellow), C6-279 (orange) and C6-280 (blue) in a medial section of the human hypothalamus. C6-278 has higher abundance values in comparison to C6-279 and C6-280. All 3 clusters map to the arcuate nucleus. **d)** Spatial mapping of AGRP clusters. Cell abundance scores for Cg-260 (yellow), C6-212 (blue) and C6-396 (orange). C6-260 (AGRP/NPY neurons) map to the ARC, C6-212 (GHRH) map to the ARC and periventricular nucleus, and C6-396 (TRH/SIM1) map to the PVN. **e)** smFISH of AGRP expression in the medial hypothalamus in a near adjacent section to the spatial transcriptomics section. No expression identified in the PVN (1) or perventricular hypothalamus (2). Abundant AGRP expression in the ARC (3). Diagram highlights where in the section each image was taken. **f)** Characterisation of MC4R populations in the human hypothalamus. (Left) Log-normalised expression of MC4R in the snRNAseq data. MC4R expression is found in several different clusters. (Right) Heatmap displaying marker genes for clusters in the 98^th^ percentile of MC4R expression and dotplot showing scaled average cell abundance of each MC4R snRNAseq cluster within each spatial transcriptomics cluster. The size of the dot represents the percentage of spots in that cluster that more than 0.05 cell abundance score for the snRNAseq population. Cluster C6-112 maps to the LPOA, C6-181 maps to the intermediate nucleus of the hypothalamus and MPOA, C6-224 shows some mapping to the MPOA/GABAergic MBH region. The rest of the MC4R clusters show low cell abundance mapping to the spatial transcriptomics dataset. Spatial transcriptomics clusters were only included in the plot if at least one snRNAseq cell type cluster mapped to a minimum of 5% of spots in a spatial transcriptomics cluster. **g)** Characterisation of MC3R populations in the human hypothalamus. (Left) Log-normalised expression of MC3R in the snRNAseq data. MC3R expression is found in several clusters, but mainly in TBX3+ region and some in the FEZF1+ region. (Right) Heat map map of cluster markers for clusters in the top 98^th^ percentile of MC3R expression, and a dot plot displaying scaled average cell abundance for each snRNAseq cluster in each spatial transcriptomics cluster. The size of the dot represents the percentage of spots in that cluster that more than 0.05 cell abundance score for the snRNAseq population. Of the 10 MC3R clusters, 9 map, at least in part, to the ARC, with cluster C6-243 mapping to the VMH.

On the other hand, when examining clusters in the top 99^th^ percentile of *AGRP* expression, we identified five clusters (**Figure 3A,B, extended data figure 6C,D,F**), four of which are located within the ARC/MBH (C6-260, C6-212 shown in **Figure 3D**, others in **Extended Data Figure 6**). The cluster with the highest level of *AGRP* C6-260 (NE_Mid_1_3_OTP_SOX5_AGRP) co-expresses *NPY* and is GABA-ergic, are likely the canonical *AGRP/NPY* neurons. Interestingly, we also detect low grade *AGRP* expression in C6-212 and C6-196 in the ARC that co-express *GHRH* and *GAL and GHSR*; and in C6-396 neurons in the PVN that co-express *AVP, CARTPT* and *TRH*, but we could not validate the presence of any *AGRP* transcripts in the PVN using spatial transcriptomics, or RNAScope (Figure 3E).

In contrast to the neuropeptidergic melanocortin neurons, the melanocortin receptors *MC4R* and *MC3R* are more diffusely expressed in the snRNAseq and spatial transcriptomics datasets (**extended figure 7**). To characterise populations expressing these receptors, we looked at clusters in the 98^th^ percentile of *MC3R* and *MC4R* expression. *MC4R* expression is found in 10 clusters, with >20% of cells expressing MC4R (**Figure 3F, Extended data figure 7A,B**). The receptor is highest expressed in C6-43 (NE_PRE_1_MYO16_CXCL14_CAV1) and is detected in 49% of the cells within the cluster. However, this cluster did not map to our spatial transcriptomics dataset (**Figure 3F; extended data figure 7A,B**). One of the *MC4R* clusters also expresses *SIM1* (C6-407), indicating they might originate from the PVN, SON or mamillary region; but similarly the spatial mapping of this cluster was not reliable. Of the top 10 *MC4R* clusters, only 3 show appreciable mapping, and these are the cholinergic cluster C6-112, the *SKOR2*-expressing cluster C6-224, and the *GLI3* expressing cluster C6-181, which are mapped to the LPOA, the medial preoptic area (MPOA), and the intermediate nucleus of the hypothalamus (IMH) respectively, in the anterior hypothalamus (**Figure 3F**).

Previously, we have shown that *MC3R* is expressed by ARC *GHRH* and *KISS1/TAC3* neurons in both mice and humans ^4^. Here, *MC3R* expression is detectable in two *GHRH* (C6-196, C6-212) and two *KISS1* (C6-288, C6-291) populations (**Figure 3G**). Spatial mapping of *MC3R* with the *GHRH* neurons (C6-196, C6-212) and the *KISS1* populations show overlapping localisation in the ARC (**Figure 3G, Extended data figure 7C,D**). We also identified a cluster of *MC3R* neurons that co-expresses *PGR* and *CALCR*, mapping near to the *KISS1* neurons in the ARC (C6-289) (**Figure 3G**). Of the top 10 *MC3R* expressing clusters, 9 mapped to the ARC, and 2 of these clusters are *POMC* clusters (**Figure 3G, Extended data figure 7**). Interestingly, we identify a population of VMH neurons that express *MC3R*: C6-243, which we have shown maps to the medial region of the VMH (**Figure 2D**).

Thus, these data provide a detailed expression profile of the human melanocortin neurocircuitry for the first time.

### Incretin Receptors in the human hypothalamus

Next, we turned our attention to the receptors of the incretin hormones GLP-1 and GIP, with both GLP1R and GIPR being targets for type 2 diabetes mellitus and obesity therapeutics ^19^ including semaglutide ^5^ and tirzepatide ^6^. *GLP1R* is expressed in various tissues including the pancreas, stomach, intestine and heart, and it is also found in the hypothalamus and the hindbrain, amongst other brain regions ^20^. There is growing evidence that GLP1 analogues such as liraglutide and semaglutide exhibit their weight-loss effects through activation of their receptors expressed in the hypothalamus and hindbrain, including by directly acting on POMC neurons in the ARC ^7^.

Defining clusters in the 98^th^ percentile of *GLP1R* expression, we identify 10 neuronal clusters. Three of the clusters (C6-414, C6-413, C6-396) express *SIM1* and *AVP* (**Figure 4A,B**). C6-414 (NE_Mid-2_SIM1_SCGN_EBF3_AVP) has the highest expression of *GLP1R* and the transcript is detected in 77.9% of the neurons within the cluster. Interestingly, we also detected co-expression of *GLP1R* and *GIPR* in 25% of cells within this cluster. All three *SIM1/AVP* clusters show overlapping mapping in the spatial data and they are mapped to the PVN and SON (C6-414 shown in **Figure 4C**, others shown in **Extended Data Figure 8A-C**). *GLP1R* is also detectable in 21.98% of *POMC/LEPR* neurons (C6-278) in the ARC (**Figure 4C**). It has been shown in the mouse hypothalamus that *Pomc/Glp1r* and *Pomc/LepR* neurons are two distinct populations in the hypothalamus ^12,21^. Of note, nearly all *POMC/GLP1R* neurons in this cluster (19.81% of whole cluster) also co-expressed *LEPR* transcripts, demonstrating differences in anorexigenic *POMC* populations between mice and human. A third population of interest is a GABAergic population of *SST* and *CALCR* co-expressing neurons (C6-209, mapping to the MBH, but outside of the ARC (**Figure 4C**). We performed orthogonal validation by single-molecule in-situ hybridisation to confirm the co-expression of *AVP+GLP1R, POMC+GLP1R, SST+CALCR+GLP1R* (**Figure 4C**). C6-285 are *TBX19*-expressing neurons demonstrating some mapping to the MBH, while C6-198 are *NKX2-4* and *TRH*-expressing neurons from the periventricular hypothalamus, both of which were shown to express *Glp1r* in mice ^12^.

**Figure 4:**
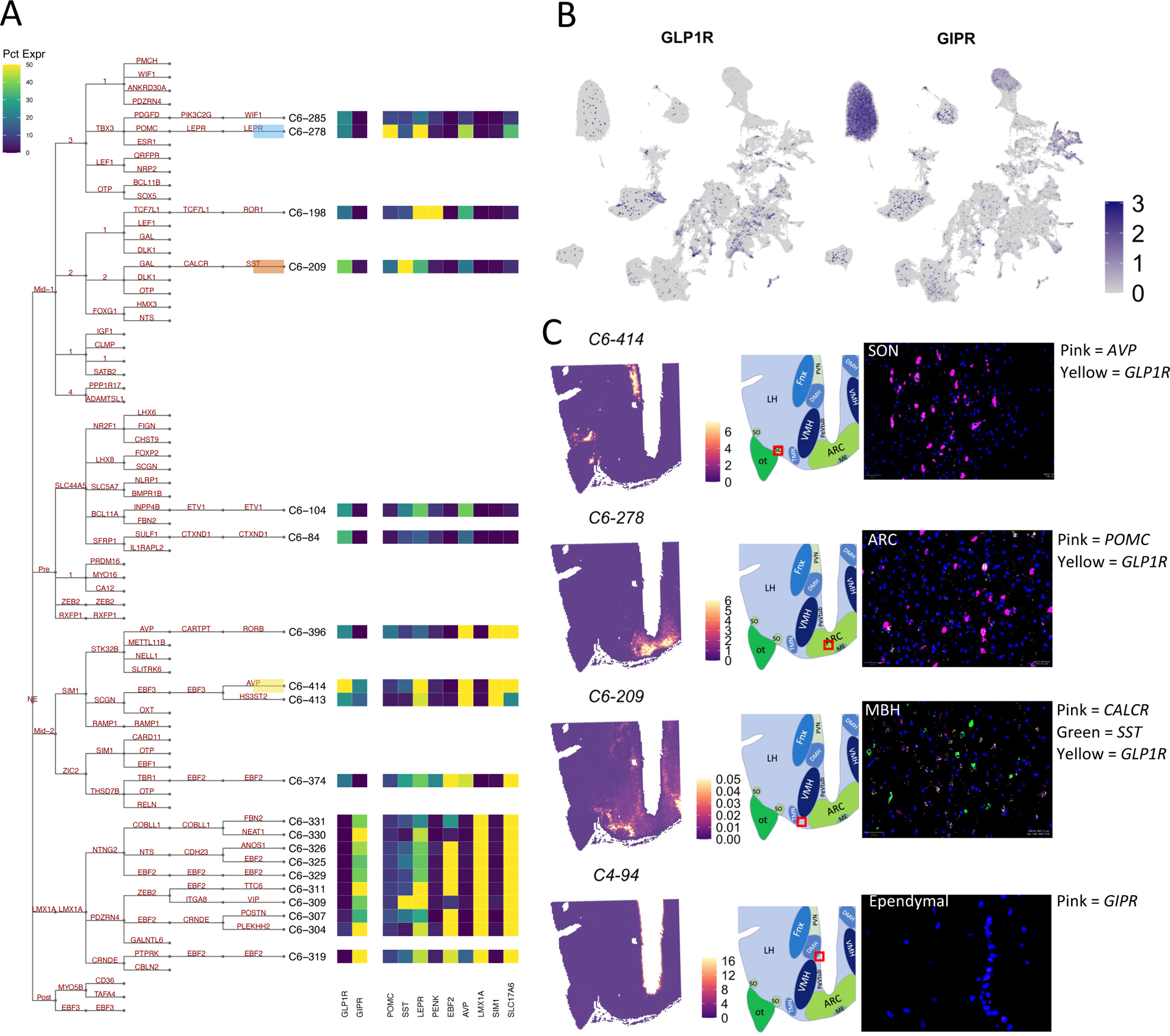
Incretin receptor expression in the human hypothalamus. **a)** Details of the branches of the clusters in the top 98 percentile of GLP1R of GIPR expression. Here, an overview of the subcluster structure is displaed and a heatmap showing the percentage expression of cells in each cluster that express each marker gene. **b)** UMAP plots showing log-normalised expression of GLP1R (left) and GIPR (right). GLP1R is found in several neuronal clusters, while GIPR is found in both neurons and non-neurons. **c)** (Left) Cell2location mapping of 3 GLP1R clusters: C6-414 (mapping to the PVN and SON), C6-278 (mapping to the ARC), and C6-209 (mapping to the MBH) and cell2location mapping of the ependymal cluster in which GIPR is expressed in. This cluster maps to the lining of the third ventricle. (Right) smFISH showing co-expression of GLP1R & AVP in the SON; GLP1R and POMC in the ARC, CALCR, SST and GLP1R in the MBH, and GIPR expression in the ependymal cell lining. Diagrams highlight where each image was taken.

In contrast, *GIPR* is expressed in both neuronal and non-neuronal populations. They are co-expressed with *LMX1A* neurons (10 clusters, **Figure 4A, Extended Data Figure 8C-E**), and interestingly, we detected *GIPR* expression in mature oligodendrocytes that map to the fornix (FX) and the optic tract (OT), and strikingly, the ependymal cells (C4-94) surrounding the 3^rd^ ventricle (**Figure 4C)**. Collectively, these data provide, for the first time, a high-resolution expression pattern of incretin hormone receptors in both neuronal and non-neuronal cells in the human hypothalamus.

### Hypothalamic neuronal populations are enriched for genes harbouring BMI-associated variants

Finally, we wanted to identify which hypothalamic cell types are implicated in the genetic regulation of obesity. To do this, we first integrated HYPOMAP with data from a common variant GWAS of body mass index (BMI) in up to 806,834 individuals ^22^, using CELL-type Expression-specific integration for Complex Traits (CELLECT) ^23^ and Multi-marker Analysis of GenoMic Annotation (MAGMA) ^24^. We found that 182 out of the 458 tested hypothalamic cell types showed significant enrichment in the BMI GWAS signals (at Bonferroni corrected P<0.05/458, **Figure 5A, Supplementary Table 15**). The majority of the enriched cell populations mapped to neurons in the middle hypothalamus (166/182), with a few mapping to the anterior (18/182), posterior (1/182) and general hypothalamus (5/182). The most significantly enriched neuronal cluster C6-179 (P=6.49Non-neuronal populations did not10-20) is an *HMX3*- and *NPSR1*-expressing cluster from the IMH. Additionally, cluster C6-236, which maps to the lateral VMH and is marked by *FEZF1*, *NTNG1* and *FAM9B* was highly enriched in the BMI GWAS (5^th^, P= 2.31×10-18; see **Extended Data Figure 4** for mapping). Notably the *SST*-, *CALCR*- and *GLP1R*-expressing cluster C6−209 shown in **Figure 4C**, also was highly enriched in the BMI GWAS (17th, P=1.28×10-14). Non-neuronal populations did not show any enrichment in the BMI GWAS. We next sought to identify ‘effector’ genes which might be driving these associations, defined as those in the 95^th^+ centile for cell-type specificity and in the top 1000 MAGMA gene associations derived from the GWAS data (*via* CELLECT GENES). This identified a total of 375 effector genes (**Supplementary Table 16**), the majority of which (362/375) were identified as effector genes in neuronal populations and (290/375) in the BMI GWAS enriched neuronal subpopulations.

**Figure 5.**
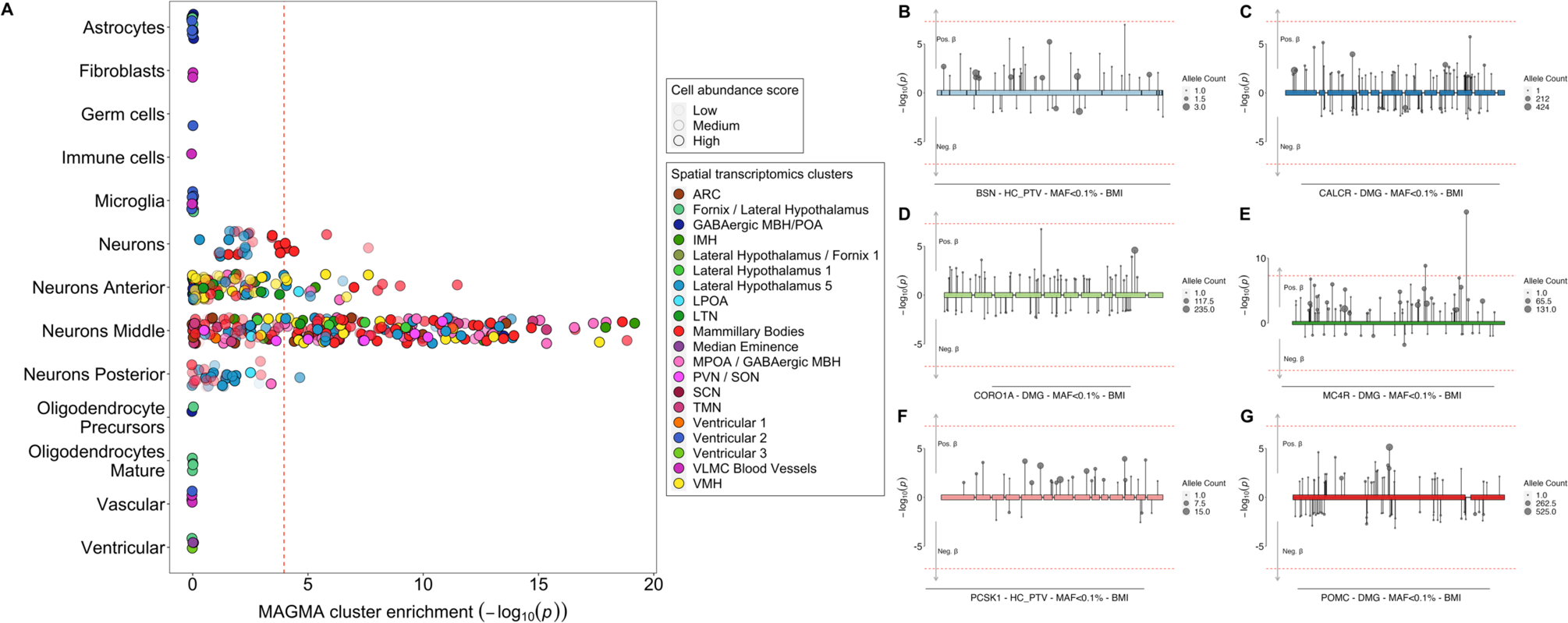
| Neuronal clusters are enriched for genes linked to BMI variation in the general population. **a)** Prioritisation of 458 human hypothalamic cell types identified 182 cell types as significantly enriched for associations in the BMI GWAS, mapping to four neuronal populations. Cell types were grouped into broader categories for visibility, as in Figure 2. The dashed line indicates a Bonferroni significance threshold, P<0.05/458. Clusters are coloured based on their regional abundance, as seen in Figure 2. Extended data are shown in Supplementary Table 15. **(b-g)** Variant-level associations in identified effector genes in UK Biobank. Rare exome variant associations with BMI for variants within BSN **(b)**, CALCR **(c)**, CORO1A **(d)**, MC4R **(e)**, PCSK1 **(f)**, POMC **(g)**. Variant collapsing masks included variants with a minor allele frequency (MAF) < 0.1% and annotated as either high-confidence protein truncating variants (HC_PTV) or HC_PTV plus missense variants with a high CADD score (>=25, denoted DMG). Each variant association is represented by a circle and vertical line: the line length indicates the P-value (−log10), in the direction of its effect on BMI in carriers of the rare allele, and the circle size indicates the number of carriers of each variant (i.e. the allele count). Exons are indicated by the boxes and connected by the intron line. Extended data are shown in Supplementary Table 16 and 17.

To determine if disruption of these effector genes influences obesity risk at the population level, we used exome-sequencing data from the UK Biobank study (n=419,692) ^25^. We performed rare-variant burden tests towards BMI for variants with a minor allele frequency <0.1% that were either protein truncating (PTVs) or missense variants with a high CADD score (≥25, see Methods). We found that carrying rare deleterious variants in 6 (of the 375) effector genes was significantly associated with changes in BMI (at P<0.05/375, **Figure 5B-G, Supplementary Table 17**). Reassuringly, these included well-established causes of monogenic obesity and previously reported associations; *MC4R* ^26–29^, *PCSK1* ^26,30^, *POMC* ^31^ and *CALCR* ^26^. However, our analysis also highlighted two new genes; *BSN*, a presynaptic protein with a role in exocytosis-mediated neurotransmitter release ^32^ that we have recently shown is associated with increased risk of severe obesity, NAFLD and T2D (Zhao, Chukanova et al, https://doi.org/10.1101/2023.06.14.23291368), and *CORO1A* (n=415 carriers, β=0.98 ±0.215, P=5.6×10^−6^), which encodes a WD repeat protein involved in cell cycle progression, signal transduction, apoptosis, and gene regulation ^33^, a gene previously unlinked with obesity.

## DISCUSSION

In functional hypothalamic research, the vast majority of ‘ground truths’ have, until recently, emerged from murine neuroanatomical and functional studies. The maturation of single cell technologies, coupled with the availability of relevant and precious donor samples, has ushered in a new era of possibilities in human brain mapping. While there are ‘whole brain’ single cell datasets emerging, from both developing ^34^ and adult humans (Siletti et al, https://doi.org/10.1101/2022.10.12.511898), here we provide a detailed, high resolution spatio-cellular map of the adult human hypothalamus.

It is the spatial element that provides a rich and novel dimension to the increasingly ubiquitous single cell data. Often ignored non-neuronal cell types serve as prime example. The snRNAseq data reveals seven different astrocyte and two different oligodendrocyte clusters, that HYPOMAP is actually able to locate onto different areas within the hypothalamus (**Extended data figure 5**). Another advantage of combining single cell and spatial data is emerging synergy of both technologies. Because the human hypothalamic blocks used for snRNAseq typically arise from hemisected brains, the median eminence tends to get lost in the process. Thus, tanycytes, which are enriched in the median eminence, are often neglected in the snRNAseq data, with only ∼100 cells identified. Yet, the expression profile of the tanycytes can be clearly mapped onto the spatial whole transcriptome data (**Extended Data Figure 5**).

Then there is the central leptin-melanocortin appetitive control pathway, whose major components were all uncovered more than 20 years ago using genetics ^1,3^, but whose mapping in the human hypothalamus we finally reveal here for the first time. We are able to resolve spatially distinct populations of *POMC* (**Figure 3A-C**), *MC4R* (**Figure 3F**) and *MC3R* (**Figure 3G**) neurons. With up to 0.3% of the general population carrying pathogenic mutations in *MC4R* ^35^ and drugs on the market now targeting this pathway, there has never been a more relevant time to increase our understanding about this pathway in the human context.

The receptors *du jour* however, at least in terms of broadest societal relevance, are the incretin hormone receptors, GLP1R and GIPR, both key targets for anti-obesity therapy development ^19^. Here we are able to confirm that *GLP1R* in human hypothalamus is almost exclusively expressed in neurons, and identify several separate populations, including one that co-expresses *POMC* and resides in the ARC, and a number of *AVP*+ clusters mapping to the PVN and SON. In contrast, *GIPR* is expressed in both non-neuronal and neuronal populations, consistent with our previous observations in both the mouse ^12^ and human hypothalamus ^36^ and the mouse hindbrain ^37^. The *GIPR* population that intrigued us the most however, was ependymal population marked by *DTHD1* (**Figure 4C**). Heterozygous loss of function mutations in GIPR are associated with lower BMI ^38^, while pharmacological studies in humans indicate that both agonism and antagonism of this receptor can augment weight loss ^39^. With recent data showing that the GLP1R/GIPR co-agonist tirzepatide might be marginally more effective than the GLP1R mono-agonist semaglutide ^40^, does this spatial localisation of GIPR to the ependymal cells hint as to why this might be the case? Could these GIPR expressing cells be increasing access of tirzepatide to the hypothalamus? While further work will be required to address these questions, our data illuminating the high resolution expression profile of hypothalamic incretin receptors in a human context is an important first step.

When we look at HYPOMAP through a genetics lens, we find a significant enrichment in expression of BMI associated genes, specifically in neurons, which is coherent with our current understanding that the large variation in body-weight is driven primarily by neuronal mechanisms. Finally, gene burden analysis of the 375 ‘effector’ genes that drove the enrichment identified 6 genes, in which rare deleterious variants was significantly associated with changes in BMI, with four of these, *MC4R*, *PCSK1*, *POMC* and *CALCR* having well-established links to body-weight regulation. It is gratifying that our approach also highlighted *BSN*, a gene we have recently shown to be linked to obesity, and *CORO1A*, an entirely new player in the regulation of energy balance, we are reporting for the first time, thus highlighting HYPOMAP as a platform for discovery.

There are, of course, limitations to our study. First, it is crucial to remember that transcriptomic data, of all types, are designed to identify what is expressed, as opposed to what is not. Second, HYPOMAP has been derived from relatively few donors (11 for the snRNAseq dataset and 5 for the spatial data, balanced in terms of male:female), limiting us from deep quantitative analyses, such as effectively comparing differences between the male and female brain. Third, these donors were of normal weight when they died, so while of interest in and of itself, the long-term value of this data, given the role of the hypothalamus in maintaining homeostasis, is as a baseline to study this brain region in states of dyshomeostasis. This will require the difficult and long-term prospective recruitment of donors suffering from relevant diseases, in our case, severe obesity.

Given our field of expertise, it is natural that we have focussed here, initially, on characterisation of the appetitive control circuitry. However, clearly this barely begins to scratch the surface of possibilities with this dataset. We hope that by making this Human HYPOMAP completely open-access, it will help illuminate human relevant neuronal populations and circuits more broadly, thus enabling the identification of novel druggable targets for treating a wide range of conditions linked to the hypothalamus, including reproductive, circadian, and metabolic disorders.

## METHODS

### Human post-mortem sample preparation

Anonymised human samples were obtained from the MRC Brain Bank Network, in line with each bank’s Research Ethics Committee approval. Subjects were approached in life for written consent for brain banking, and all tissue donations were collected and stored following legal and ethical guidelines.

For snRNAseq, frozen blocks of post-mortem hypothalamus were sourced from adult donors with BMI ranging from 18 to 28 and no significant neuropathology. Dissections were performed following delineation of relevant anatomy in Cresyl Violet stained sections from the anterior and posterior surfaces of each sample by a consultant pathologist. Samples from the relevant region were then acquired using a punch biopsy, or macrodissected from 100um thick frozen cryostat sections spanning the whole specimen.

For spatial transcriptomics, post-mortem paraffin-embedded (FFPE) human brain samples covering the hypothalamus was obtained from the Edinburgh Brain Bank (BBN001.37136, BBN001.37137, BBN001.37298, BBN001.37298, BBN001.37242). Selection of samples and areas to include in ST analyses were based on anatomical landmarks using Luxol Fast Blue/Hematoxylin-Eosin staining of myelinated fibers and cell bodies. n=7 samples from n=5 different donors (2 males, 3 females), ranging from 50-95 years were included in the ST analyses. The BMI varied between 16-41 at the time of death and the post-mortem interval ranged from 12-102 hours.

### Nucleus dissociation and RNA-sequencing

Nuclei were isolated by dounce homogenisation and purified using a modified protocol from ^12^. Briefly, chopped samples were transferred to a 15 ml Dounce Homogenizer with 5ml homogenisation buffer (100 μM of DTT [Sigma–Aldrich, St. Louis, MO, USA], 0.1% Triton X-100 [Sigma–Aldrich], 2X EDTA Protease Inhibitor [Roche, Basel, Switzerland], 0.4 U/μl of RNasin RNase inhibitor [Promega, Madison, WI, USA, 10000 U, 40 U/ml], and 0.2 U/μl of Superase.In RNase Inhibitor [Ambion, Austin, TX, USA, 10000 U, 20 U/μl] in nuclei isolation medium [250 mM of sucrose, 25 mM of KCl (Ambion), 5 mM of MgCl2 (Ambion), and 10 mM of Tris buffer at a pH of 8.0 (Ambion) in nuclease-free water (Ambion)] with 1 μl/ml of DRAQ5 [Biostatus, Loughborough, UK]), and mechanical dissociation was performed using 10 strokes with pestle A, and 20 strokes with pestle B. Homogenates were filtered through a 100um filter and centrifuged at 600 x *g* for 5 minutes in a precooled centrifuge. Supernatant was discarded and pellet was resuspended in 27% Optiprep solution diluted in homogenization buffer and centrifuged at 13,600 x *g* for 20 minutes at 4°C. The nuclei pellet was collected and resuspended in wash buffer (1% BSA, 0.4 U/μl of RNasin, and 0.2 of U/μl of Superase.In in PBS [Sigma–Aldrich]) and centrifuged at 700 x *g* for 5 minutes at 4°C. This was repeated twice before being passed through a 40um cell strainer and this final sample was used to create sequencing libraries. For two donors, single nuclear suspensions were sorted using fluorescent-activated nucleus-sorting (FANS) on a BD FACSMelody instrument. The gating was set according to FSC, SSC and fluorescence at 647/670nm to detect DraQ5 nuclear staining, and 567nm to detect NeuN-PE staining. NeuN+ events were sorted into a collection tube to enrich for neuronal nuclei

Sequencing libraries were generated using 10X Genomics Chromium Single-Cell 3’ Reagent kits (Version 3.1) according to the standardised protocol. cDNA was amplified for 19 cycles. Paired-end sequencing was performed using an Illumina NovaSeq 6000.

### Sequence alignment, cell calling and QC

Raw sequence reads were mapped and genes counted based on the Human GRCh38, Ensembl 98 gene model, both using 10X Genomics CellRanger V5-6 (https://support.10xgenomics.com/single-cell-gene-expression/software/pipelines/latest/what-is-cell-ranger) using the following parameters (-- include-introns). Cellbender 2.0 ^41^ was used to recalibrate UMI counts and cell calling.

After removal of flagged nuclei, our nucSeq dataset included 571,091 from 58 samples, that contributed between 748 and 45,771 cells. We used scran‘s quickCluster function ^42^ to obtain an intial set of clusters that were used as input cluster assignments to scDblFinder which was run with multiSampleMode set to “split” ^43^. We additionally ran an initial Seurat based processing of the whole dataset, including detection of highly variable features, scaling of data, PCA and preliminary clustering ^44^. All cells detected by scDblFinder as Doublets or cells that were part of seurat clusters with more than 75% of Doublets cells were removed. We further filtered the data using the sample-based thresholds and additionally set a global threshold of maximum mitochondrial RNA of 10% and a minimum of 800 UMIs per cell. After filtering the dataset for Doublets and low quality cells, it included 353,678 cells from the 58 samples, that contributed between 609 and 20,424 cells.

We extracted the processed nucSeq data of all hypothalamus samples (ROIGroupCoarse = “Hypothalamus”) from the loom file published by Siletti et al (https://doi.org/10.1101/2022.10.12.511898). This included a total of 134,471 cells that we merged with data from our own study.

### snRNAseq Integration

Our combined human dataset includes 82 10x samples from 11 different donors and two independent studies with a total 488,149 cells after merging and initial quality control. To integrate all cells and make the data comparable we used scvi-tools ^45^, which we have previously shown to be a powerful integration tool that preserves cell type purity while removing batch differences ^12^. Similar to our previous study we optimized the main hyperparameters of scvi by running a grid search over pre-defined parameter ranges using our published pipeline (https://github.com/lsteuernagel/scIntegration). scIntegration evaluates different scvi model outputs for mixing of samples (using the entropy of the sample distribution in each cell’s nearest neighbors), the purity of cells (cell type distribution in each cell’s nearest neighbors) and the average silhouette width for cluster separation. We defined a set of ground truth cell types using signatures for mouse glial cell types from our mouse HypoMap and additionally added a set of manually curated neuron signatures. We then visualized the hyperparameters of all runs by the evaluation metrics to choose a final set of optimal parameters (Method Figure 1b-d). Overall, all models integrated the data well and we mostly found small improvements. The final scvi model was trained for 100 epochs with a dropout rate of 0.1. The model had 2 layers and 256 nodes per layer (n hidden) and the latent space had 80 dimensions. All other parameters were set to default.

### snRNAseq clustering and annotation

The low dimensional latent space from the final scvi model was used for downstream analysis. We adapted our previous dataset harmonization pipeline^12^ (https://github.com/lsteuernagel/scHarmonization) for many of the following steps, but changed it where necessary. We started with an initial round of clustering and annotated these clusters using marker gene signatures for major cell types, including some extra-hypothalamic ones (Method Figure 1a). We found multiple clusters of cells that likely reside outside the hypothalamus (e.g., SCL17A7+ neurons or thalamic SHOX2 neurons). After annotating all cells, we removed the likely extra-hypothalamic clusters and some other clusters representing low-quality cells leaving us with a final dataset of 433,369 cells. Due to the imbalance of major cell type distribution (e.g., 40.4% of all cells are oligodendrocytes) we split the data into 4 main subsets for clustering and tree building: neurons, oligodendrocytes, astrocytes and other non-neuronal cells. For each dataset multiple flat leiden clusterings were combined into a consensus subtree using mrtree ^46^ and marker genes of each cluster vs all others as well as only its sibling nodes within the subtree were calculated using a batch-stratified Wilcoxon rank sum test. The subtrees were pruned by merging nodes with insufficient differences (less than 15 marker genes). We then merged all 4 subtrees into the final clustering tree which spans 7 different levels (C0-C6) with 4 to 458 distinct clusters, however, for non-neuronal cell types only up to 5 levels exist. For each node in the tree the most informative marker gene was selected based on global differences to all other clusters and on local differences to its sibling clusters ^12^. These marker genes are shown on the edges of the circular tree. By concatenating genes from subsequent levels, a full cluster annotation can be constructed. The informative marker genes are used to name the clusters, but are not necessarily exclusive marker genes. Especially on higher tree levels it was not possible to find optimal single marker genes, hence we manually renamed many clusters with their commonly used names or a simple numbering scheme.

### 10X Genomics Visium CytAssist Spatial Transcriptomics

5um FFPE sections were prepared using a microtome (Leica) in an RNase free environment and mounted onto positively charged slides. The sections were then stored at room temperature until use. Slides were processed for spatial transcriptomics according to the 10x Genomics Visium CytAssist Version 2 protocols. Briefly, samples were deparaffinized in Xylene, a series of concentrations of Ethanol solutions (100% - 70%) and immersed in water prior to Haematoxylin and Eosin staining (H&E). Once stained, samples were cover slipped using a glycerol mountant and images using a VS200 slide scanner (Olympus Life Science) at 20X magnification (air objective, 0.8 NA). Coverslips were removed and samples underwent destaining, decrosslinking and were incubated overnight with the 10x Genomics Visium Human WT Probes version 2 (Pleasanton, CA, USA). Following this, slides were loaded at the appropriate orientation, along with the Visium 11×11mm gene expression slide, onto a CytAssist (10xGenomics), where hybridized probes were released from the tissue and ligated to spatially barcoded oligonucleotides on the Visium Gene expression slide. A tissue image was taken on the CytAssist at 10X magnification, for downstream alignment of library to the tissue section. Barcoded ligation products were then amplified to create a cDNA library for sequencing.

Libraries from the seven samples were pooled and sequenced on a NovaSeq 6000 sequencing platform (Illumina), using a NovaSeq 6000 S2 Reagent Kit v1.5 (Illumina) according to the manufacturers instructions. Subsequently, fastq files were generated for each sample, reads were aligned to their corresponding probe-sequences (Visium human transcriptome probe set v2, based on GRCh38 2020-A) and mapped back to the Visium spot where a given probe was originally captured, and finally aligned to the original HE-stained image of the tissue section using SpaceRanger version 2.0.0 (10X Genomics).

### Spatial transcriptomic data analysis

Across the seven samples, the average median number of paired-end reads per Visium spot equalled 37,863, with the min-max range from 29011 to 48778. The average median number of detected genes per spot equalled 3982 (min-max range of median values: 1405 to 7051). The average sequencing saturation equalled 0.65. Furthermore, for each individual sample, graphs with a) sequencing saturation and b) detected number of genes plotted as a function of median number of reads per spot revealed the plateau phase was either obtained or clearly approached, i.e., very little benefit would be gained from even deeper sequencing.

### Spatial Transcriptomics data pre-processing

The number of genes/spot and counts/spot was inspected for each tissue section individually using the Loupe browser to identify whether there were areas of the sample that had unusually low/high counts that are likely artefacts from the experimental procedures. These spots were identified and removed from downstream analysis.

For visualisation of gene expression in the spatial transcriptomics data, data was analysed using Seurat (version 4.3.0) ^47^. Raw count matrices along with spatial barcode coordinates for each sample were loaded, and data was log-normalised for visualisation of transcript expression.

### Integration of snRNA-seq and ST data: Cell2Location

We utilised the Cell2Location tool (Version 0.1.3) ^15^ to predict the locations of snRNA-seq cell populations in the ST data. We utilised the entire snRNA-seq dataset as a reference, and estimated reference cell type signatures for clustering levels C1:C6. We included genes that were expressed in at least 8% of cells, and genes expressed in at least 0.05% of cells, if the non-zero mean was greater than 1.4. We estimated reference signatures using the negative binomial regression model, accounting for the effects of donor, sex, batch, and dataset.

For each cluster level, we trained the Cell2Location model with a detection alpha of 20, and 3 cells per location as hyperparameters, and trained for 30,000 epochs, with the final gene list including genes expressed in both the snRNA-seq and ST dataset. Results were visualized using scanpy and Seurat. The plots represent the estimated abundance of cell types at each location.

To cluster the spatial transcriptomics spots, we utilised the cell abundance matrix which had snRNAseq clusters as columns, and spot barcodes as rows with cell abundance scores in each cell. Using this we used K nearest neighbours and Leiden clustering to cluster the spots according to the cell abundance scores. Each cluster was then labelled based on the hypothalamic region in which most spots appeared. If more than 1 region was covered, then the top two regions were used to label the cluster.

### Fixation & parafinization of human samples

Three independent human samples were used to assess ependymal and tanycyte expression markers. Fresh post-mortem human hypothalamus 2 x 3 x 1 cm blocks (<24h post-mortem) were incubated for 16h in 10% NBF and then further fixed for 48-72h in 4% PFA. Brain blocks were dehydrated in a series of ethanol treatments (70% (16h, 2×4h), 80% (16h, 2×4h), 96% (16h, 2×4h), 100% (16h, 1×4h)). The blocks were then incubated for 3.5 days in Xylol, followed by 2 incubations in fresh paraffin (5h, 16h) until the blocks were poured into forms. Brain blocks were sliced (5 µm) and mounted on Superfrost (Thermo Fisher) glass slides and stored at RT.

#### Donor info

91y, female, BMI:15.4, ICD codes for diseases: E41, E86, E48, I10.9, R54 (malnutrition, dehydration, hypertension, age related physical debility)

90y, female, BMI: 24.6, ICD codes for diseases: N17, E86, F03, I10, E11 (acute kidney failure, dehydration, unspecified dementia, primary hypertension, type II diabetes)

67y male, BMI 25.5, ICD codes: K70.4, K70.3, D64.9, J96.9 (alcoholic liver disease, anemia, respiratory failure)

### smFISH

5um FFPE sections were cut and mounted onto positively charged slides. Multiplex fluorescence RNAscope (ACDBio) was performed using a Bond RX fully automated research stainer (Leica), the RNAscope LS multiplex fluorescent reagent kit (Advanced Cell Diagnostics, Bio-Techne) and probes specific for *GLP1R* (Cat# 519828), *GIPR* (Cat# 471348), *SST* (Cat# 310598), *POMC* (Cat# 429908), *AVP*(Cat# 401368) (Advanced Cell Diagnostics, Bio-Techne). Slides were baked and deparaffinized prior to heat-induced epitope retrieval at 95°C for 30 minutes using Bond ER Solution 2. Next ACD Enzyme (ACDBio) was added, and slides were incubated at 40°C for 15 minutes. Hybridization, amplification, and detection was performed according to the ACD Multiplex Protocol P1. Final detection was achieved with the Opal-570 and Opal-690 fluorophore reagent packs (Akoya BioSciences, Inc.,diluted 1:1000), and samples were conterstained with DAPI (ACD) to mark cell nuclei and coverslipped with ProLong Diamond antifade mountant (ThermoFisher Scientific) before imaged using the VS200 slide scanner (Olympus Life Science) at 20X magnification (air objective, 0.8 NA).

### Cell type enrichment & BMI associations

Cell type specificity matrices were generated using CELLEX software v1.2.2 ^23^. Due to memory limits, we performed bootstrapping by sampling the HYPOMAP dataset randomly into 9 smaller datasets, each containing 100K cells. CELLEX was then performed on each of the subsets, and the mean values were taken forward for the subsequent enrichment analysis.

Using the resulting cell-type specificity matrices, we ran CELL-type Expression-specific integration for Complex Traits (CELLECT) ^23^ with Multi-marker Analysis of GenoMic Annotation (MAGMA) ^24^, alongside GWAS data from the GIANT BMI meta-analysis (Nmax=806,834) ^22^, to prioritise hypothalamic cell types that showed enrichment in the BMI GWAS. CELLECT-MAGMA (version 1.3.0) was run with default parameters across the 458 tested hypothalamic cell types, setting the multiple-test corrected significance threshold at P<0.05/458 and followed-up by CELLECT-GENES, but setting the percentile cutoff to 95. CELLECT-MAGMA was also run on the above mentioned subsets as a sensitivity analysis (Supplementary Figure 9).

Exome sequencing-based rare variant burden analyses, as described in Gardner et al. ^48^ using data from up to 454,787 individuals from the UK Biobank study ^25^ via the UK Biobank Research Access Platform (https://ukbiobank.dnanexus.com). Variants were then annotated with the ENSEMBL Variant Effect Predictor (VEP) ^49^ v10448 with the ‘everything’ flag and the LOFTEE plugin ^50^ and prioritised a single MANE v0.97 or VEP canonical ENSEMBL transcript and most damaging consequence as defined by VEP defaults. To define Protein Truncating Variants (PTVs), we grouped high-confidence (as defined by LOFTEE) stop gained, splice donor/acceptor, and frameshift consequences. All variants were subsequently annotated using CADD (v1.650) ^51^. BMI for all participants was obtained from the UK Biobank data showcase (field 21001). After excluding individuals with missing data, 419,692 individuals with BMI measures remained for downstream analysis. To assess the association between rare variant burden and BMI we implemented BOLT-LMM (v2.3.551) ^52^, using a set of dummy genotypes representing the per-gene carrier status. For the latter, we collapsed variants with a minor allele frequency (MAF) < 0.1% across each gene and defined carriers of variants as those with a qualifying high confidence PTV (HC PTV) as defined by VEP and LOFTEE or “Damaging” variants (DMG), including missense variants with a CADD score >=25 and the aforementioned HC PTVs. Genes with fewer than 10 carriers were excluded. BOLT-LMM was run with default settings and the ‘lmmInfOnly’ flag and all analyses were controlled for sex, age, age2, WES batch, and the first ten genetic ancestral principal components as calculated ^25^. Gene-level BOLT association summary statistics were then extracted for the 375 identified effector genes, setting the multiple-test corrected threshold at P<0.05/375.

Finally, to identify which GWAS signals were proximal to the identified effector genes, we also performed signal selection on the GIANT BMI GWAS meta-analysis. GWAS summary statistics were filtered to retain variants with a MAF>0.1% and which were present in at least half the contributing studies. Quasi-independent genome-wide significant (P<5×10-8) signals were initially selected in 1Mb windows and secondary signals within these loci were further selected via conditional analysis in GCTA ^53^, using an LD reference derived from the UK Biobank study. Primary signals were then supplemented with unlinked (R2<5%) secondary signals, whose association statistics did not overtly change in the conditional models. Signals were mapped to proximal effector genes, within 500kb windows. For genes within 500kb of multiple GWAS signals, the most significant signal is shown in Supplementary Table 17.

Results from CELLECT and exome associations were visualised using ggplot2 (version 3.4.2) in R (version 4.2.1).

## Extended Figure Legends

**Extended Figure 1:** snRNAseq data integration **a-c)** Evaluation metrics for scvi integration results of the combined dataset (see methods for a brief explanation of metrics) used to select the final set of hyperparameters. Boxplots show evaluation metrics of different hyperparameters runs colored by the number of layers in the model and stratified by the highly variable feature set size (a), the number of training epochs (b) and the number of latent space dimensions (c). **d)** Scatter plot of purity vs mixing metric colored by average silhouette width of all runs. PCA alone achieved good purity but did not integrate the data as well as scvi. **e)** UMAP plot of the dataset after integration but before removal of additional cell types such as non-hypothalamic neurons.

**Extended Figure 2: snRNAseq reference atlas of the human hypothalamus**. **a)** UMAP plot of the integrated atlas colored by the two contributing studies. Overall, the data is well mixed, but the contribution to different types of cells differs, with Siletti et al. (121,405) contributing more cells to many of the neuronal clusters and the newly generated data (311,964 cells) containing more non-neuronal cells. **b)** UMAP plot of the integrated atlas colored by clustering level C3. **c)** Dotplot of informative marker genes used for annotation on level C3 as shown in (b). If no informative marker gene was used to annotate the cluster, the next best marker gene was included. Dot size corresponds to the percentage of expressing cells in that cluster and color intensity to the average expression level. The dotplot shows that although all many the marker genes are strongly expressed in their respective clusters, they are not necessarily exclusive and do not always reach very high percentage of expression.

**Extended Figure 3: Key transcription factors in the snRNAseq reference atlas of the human hypothalamus and in the 7 spatial transcriptomics sections**. **a-f)** UMAP plots showing per cell expression, and spatial expression plots showing per spot expression of the transcription factors *MEIS2, LHX6, FEZF1, TBX3, SIX3, OTP, SIM1 and FOXB1*.

**Extended Figure 4: Spatial Mapping of 3 branches from C3-18 NE_Mid-1_4_ADAMTSL1 at C6 level in the hypothalamus using Cell2Location.**

Cell2location mapping of 16 C6 clusters from the *C4-51:* NE_Mid-1_4_ADAMTSL1_WDR49, C4-50: NE_Mid-1_4_ADAMTSL1_ITGA8, C4-49: NE_Mid-1_4_ADAMTSL1_FEZF1 branches in a section of the human hypothalamus. Each cluster maps to spatially distinct areas of the ventromedial hypothalamus.

**Extended Figure 5: Non-neuronal cell types in the human hypothalamus.**

**a)** Spatial mapping of astrocyte populations at the C3 cluster level. (Left) subset of the astrocyte branches from the mrtree. (rest): Cell2Location mapping of each of the 6 astrocyte C3 clusters, demonstrating spatially distinct astrocytic populations. **b)** Spatial mapping of oligodendrocyte populations as the C2 level. (Left) subset of the oligodendrocyte branches from the mrtree. At C2 level, mature oligodendrocytes split into two main clusters, marked by differential expression of *OPALIN* and *ACSBG1.* Cell2Location maps these two main populations to distinct locations in the hypothalamus, and example of this shown in the right plot. In the most anterior section of the hypothalamus, *OPALIN+* oligodendrocytes map to the anterior commissure, and ACSBG1+ oligodendrocytes map to the optic chiasm, with both cell types mapping to the lateral preoptic area. **c)** Mapping of ependymal cell types in the hypothalamus. (Left) A subset of the ventricular branches from the mrtree. (Right) spatial mapping of tanycytes and DTHD1+ ependymal cell types in the hypothalamus. Tanycytes show distinct mapping to the median eminence and the base of the third ventricle, whereas ependymal cells map to the lining of the third ventricle. **d)** Log-normalized expression of *CRYM* and *FRZB,* both previously identified to be markers of tanycyte subtypes, in the third section below the third ventricle. (Below) Log-normalized expression of *DIO2*, *FZD5*, *STOML3* and *LPAR3,* marker genes for tanycytes and/or ependymal cells with complementary RNAscope.

**Extended figure 6: Leptin-melanocortin mapping: POMC and AGRP**

**a)** Spatial expression of POMC (log-normalized per spot value) on each of the 7 human hypothalamus sections. **b)** table showing information of top POMC expressing clusters including the % cells expressing POMC, average POMC expression, marker genes, and the top region cluster of the hypothalamus each cluster maps to. **c)** Spatial expression of AGRP (log-normalized per spot value) on each of the 7 human hypothalamus sections. **d)** table showing information of top AGRP expressing clusters including the % cells expressing AGRP, average AGRP expression, marker genes, and the top cluster of the hypothalamus each cluster maps to. Dot plot showing scaled average cell abundance of clusters in the top 99 percentile of **e)** POMC and **f)** AGRP expression in the snRNAseq data. Dot size represents the percentage of spots in each cluster that had a cell abundance score greater than 0.05. 5 of the 6 POMC clusters mapped to the ARC C6-223 showing low abundance throughout the ST data. 4 or the 5 AGRP clusters mapped at varying abundance levels to the ARC, with cluster C6-396 mapping to the PVN.

**Extended figure 7: Leptin-melanocortin mapping: MC4R and MC3R**

**a)** Spatial expression of MC4R (log-normalized per spot value) on each of the 7 human hypothalamus sections. **b)** table showing information of top MC4R expressing clusters including the % cells expressing MC4R, average MC4R expression, marker genes, and which region of the hypothalamus each cluster maps to. **c)** Spatial expression of MC3R (log-normalized per spot value) on each of the 7 human hypothalamus sections. **d)** table showing information of top MC3R expressing clusters including the % cells expressing MC3R, average MC3R expression, marker genes, and which region of the hypothalamus each cluster maps to.

**Extended figure 8: Incretin receptors in the human hypothalamus.**

**a)** Spatial expression of GLP1R (log-normalized per spot value) on each of the 7 human hypothalamus sections. **b)** table showing information of top GLP1R expressing clusters including the % cells expressing GLP1R, average GLP1R expression, marker genes, and which region of the hypothalamus each cluster maps to. **c)** Scaled average cell abundance of GLP1R clusters (left) and GIPR clusters (right). Dot size represents the percentage of spots in each spatial cluster that has a cell abundance greater than 0.05. Of the GLP1R clusters, 3 mapped to the PVN and SON, and 1 mapped to the ARC. Others show some low mapping to other regions of the hypothalamus. Of the GIPR clusters 1 mapped to the mamillary body region and another showed some mapping to the lateral hypothalamus. Spatial clusters were included in the plot if at least 5% of spots displayed mapping of one of the snRNAseq clusters. **d)** Log-normalised spatial expression of GIPR on each of the 7 human hypothalamus sections. **e)** table showing information of the top GIPR expressing clusters including the % cells expressing GIPR, average GIPR expression, marker genes and which region of the hypothalamus each cluster maps to.

**Extended Figure 9 | Correlation of MAGMA enrichment across different subsets of cell populations.** Cell-type prioritisation across each the 458 human hypothalamic cell types, was compared between three different subsets of 100K cells and also enrichment calculated using reference signature values generated from Cell2Location (on the Y axes), and the dataset used in discovery (on the X axes). The Pearson correlation for each comparison is displayed.

### Acknowledgements

The authors thank Jeanette Bannebjerg Johansen, Heidi Solvang Nielsen and Jeanette Juul from Novo Nordisk for technical support on human brain histology and spatial transcriptomics. Authors also acknowledge The MRC brain network, The Edinburgh Brain and Tissue Bank for providing human brain tissue samples. We would like to thank Professor Colin Smith from The Edinburgh Brain and Tissue Bank, MRC London Brain Bank for Neurogenerative Diseases and Funder, the Cambridge Brain Bank (supported by the NIHR Cambridge Biomedical Research Centre). We would also like to thank the South West Dementia Brain Bank (SWDBB) for providing brain tissue for this study. The SWDBB is part of the Brains for Dementia Research programme, jointly funded by Alzheimer’s Research UK and Alzheimer’s Society and is supported by BRACE (Bristol Research into Alzheimer’s and Care of the Elderly) and the Medical Research Council. Tissue samples and associated clinical and neuropathological data were also supplied by the Parkinson’s UK Brain Bank, funded by Parkinson’s UK, a charity registered in England and Wales (258197) and in Scotland (SC037554). We also wish to thank Prof. Ingo Bechmann (University of Leipzig/Gremany) for access to human brain samples employed for the the in situ hybridization experiments. This research has been conducted using the UK Biobank Resource under Application Number 9905.

## Funding

JT, KR, BYHL and GSHY are supported by BBSRC Project Grant (BB/S017593/1) and the MRC Metabolic Diseases Unit (MC_UU_00014/1). GKCD is supported by a BBSRC iCASE studentship co-funded by Novo Nordisk. KAK and JRBP are supported by the UK Medical Research Council (Unit programmes: MC_UU_00006/1 and MC_UU_00006/2). Next-generation sequencing was performed at the Wellcome–MRC IMS Genomics and Bioinformatics Core supported by the MRC (MC_UU_00014/5) and the Wellcome (208363/Z/17/Z) and the Cancer Research UK Cambridge Institute Genomics Core. These funding sources had no role in the design, conduct, or analysis of the study or the decision to submit the manuscript for publication.

## Competing interests

SL, JPW, LBK and CP are Novo Nordisk employees and/or shareholders. GSHY receives grant funding from Novo Nordisk and consults for both Novo Nordisk and Eli Lilly.

