## extended data figures for "Human HYPOMAP: A comprehensive spatio-cellular map of the human hypothalamus"

### Extended data Figure 1: snRNAseq data integration

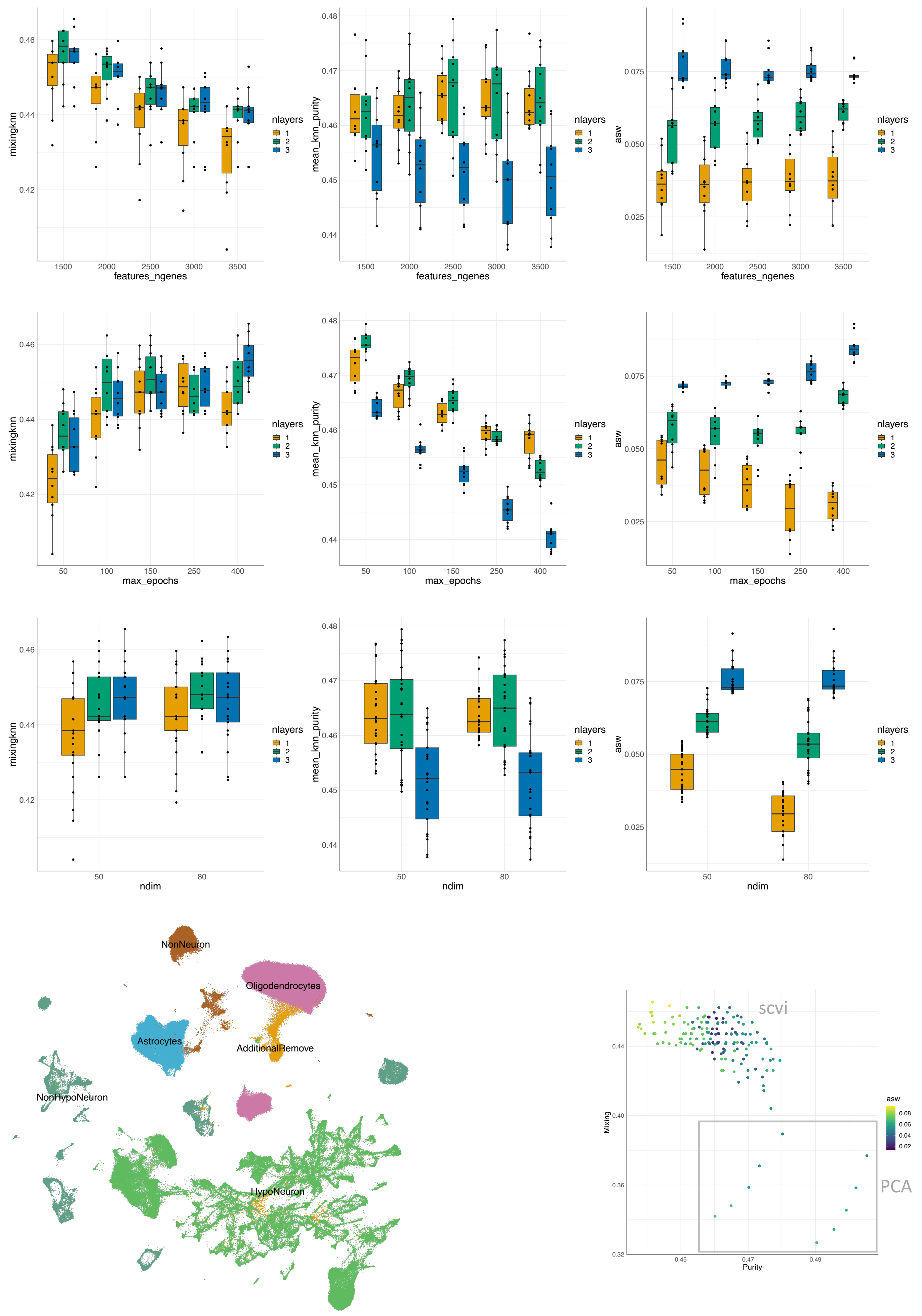

### Extended data Figure 2: snRNAseq dataset distribution and cluster markers on level C3

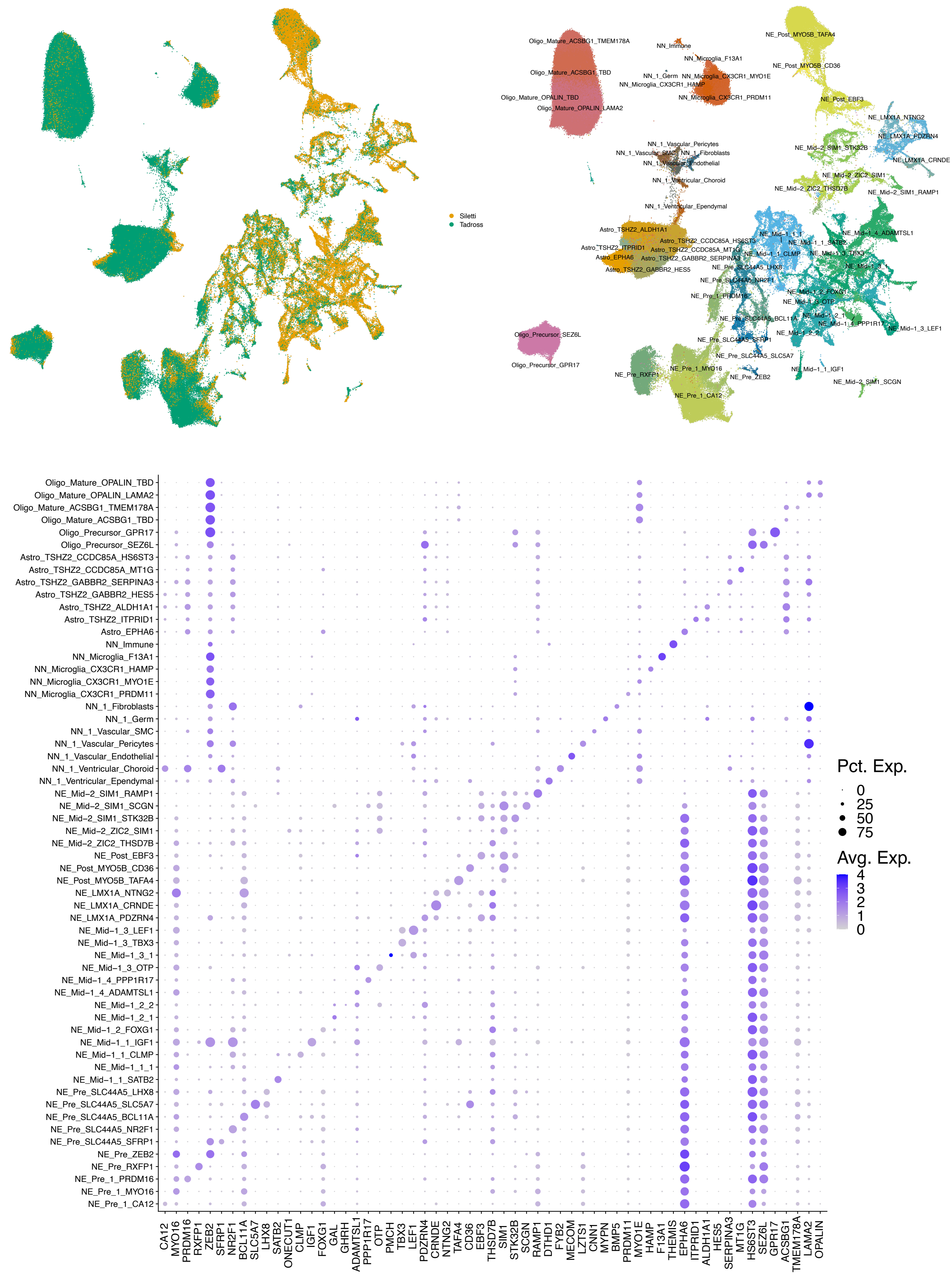

Extended data Figure 3: snRNAseq transcription factor expression levels

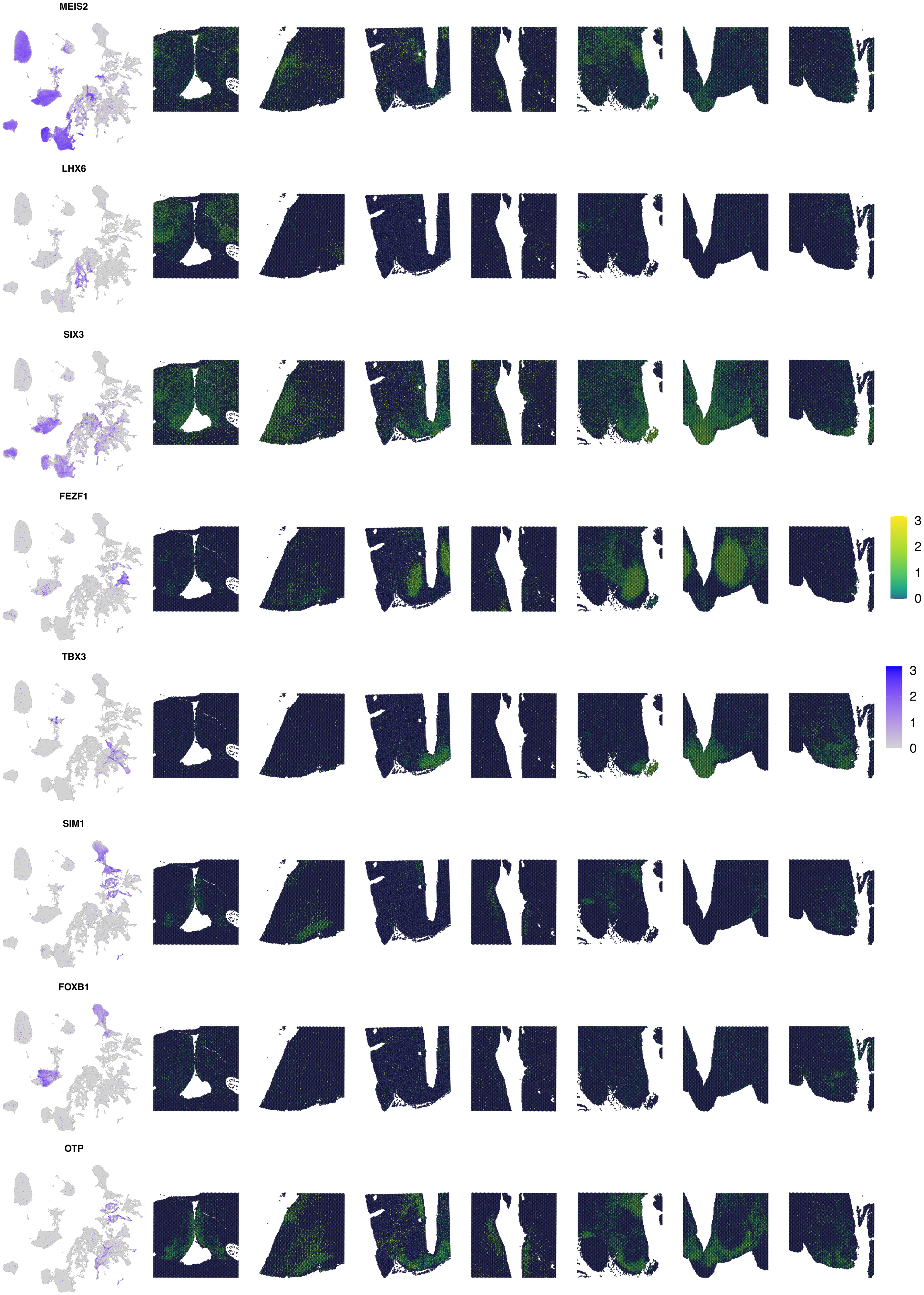

### Ext 4: Spatial mapping of ADAMTSL1 branches

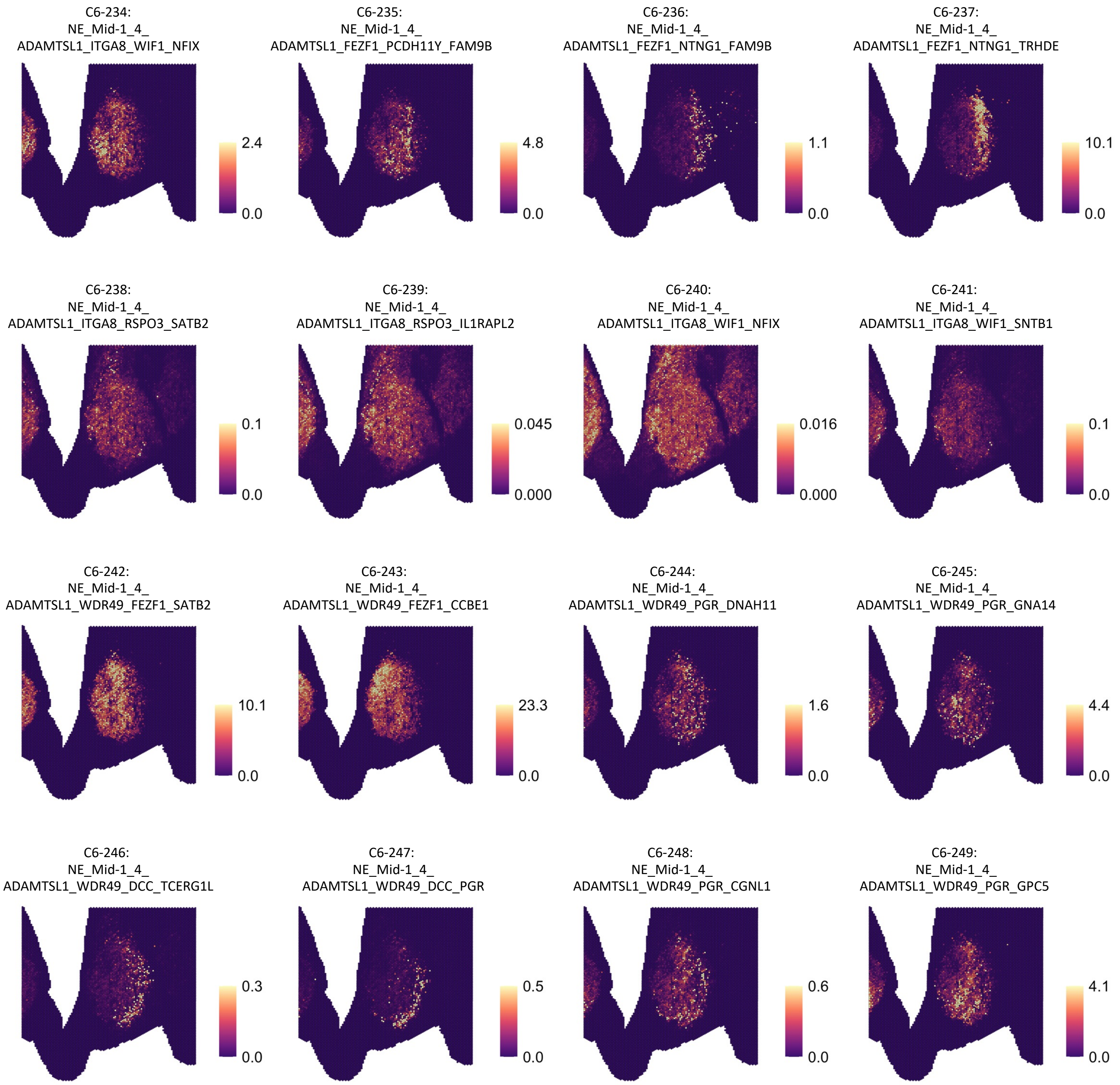

### Ext 5: Non neuronal clusters

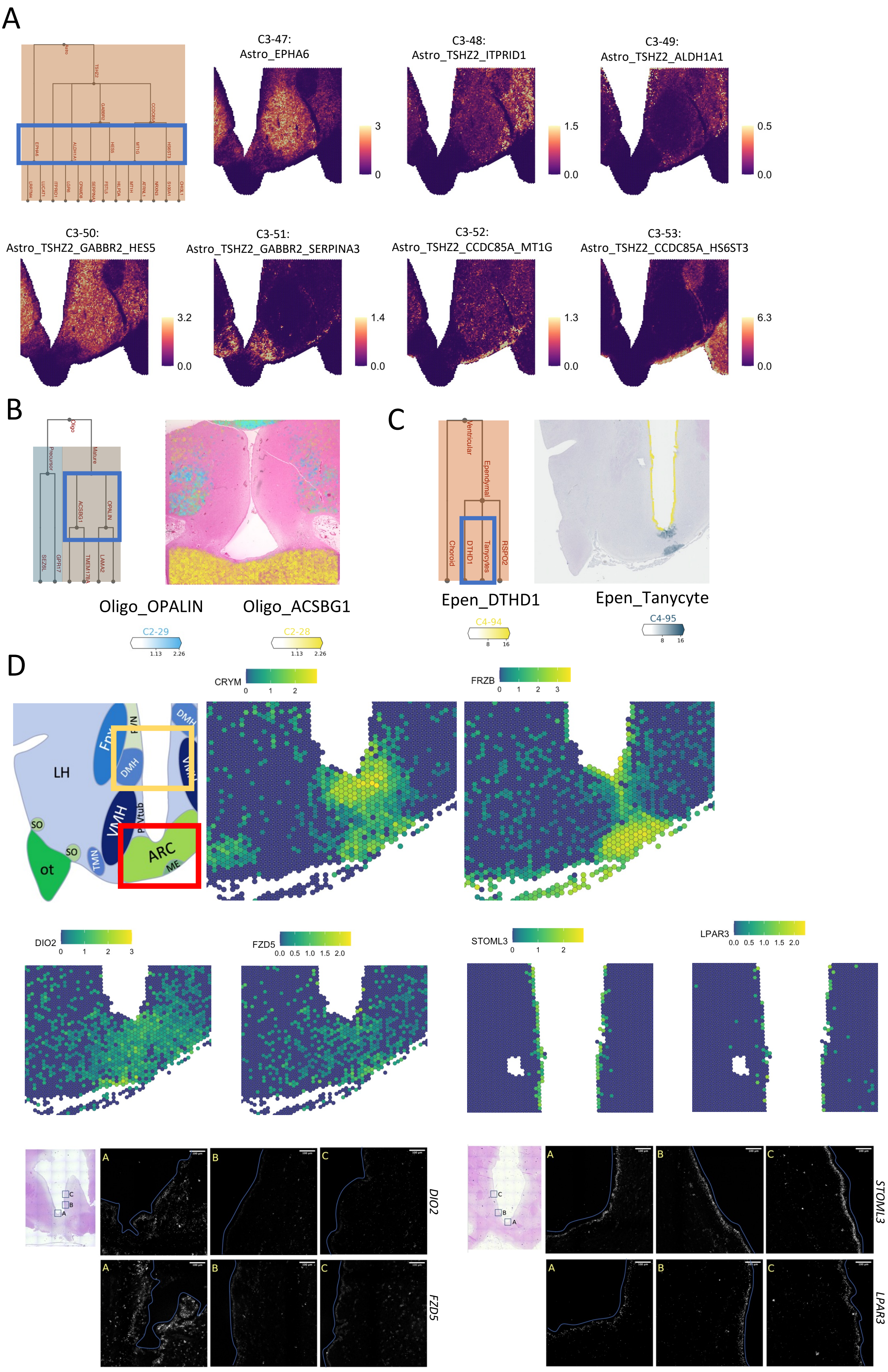

### Extended data 6: Mapping POMC & AGRP populations

A

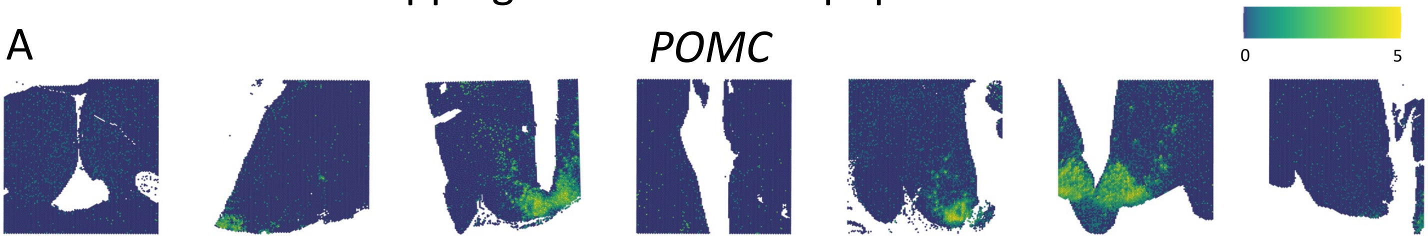


B

| Cluster | Cluster Name | Number of Cells | Number of POMC+ cells | % POMC+ | Average Expression | Cluster Marker Genes | Spatial Mapping |
|---|---|---|---|---|---|---|---|
| C6-278 | NE_Mid-1_3_TBX3_POMC_LEPR | 414 | 339 | 81.88 | 29.50 | POMC\|WIF1\|GABRE\|TBX3 | Arcuate Nucleus |
| C6-279 | NE_Mid-1_3_TBX3_POMC_CALCR | 351 | 196 | 55.84 | 17.25 | CALCR\|POMC\|WIF1\|PGR | Arcuate Nucleus |
| C6-280 | NE_Mid-1_3_TBX3_POMC_ANKRD30A | 384 | 156 | 40.63 | 6.34 | SOX3\|PGR\|POMC\|TBX3 | Arcuate Nucleus |
| C6-289 | NE_Mid-1_3_TBX3_ESR1_CALCR | 143 | 53 | 37.06 | 0.55 | PGR\|CALCR\|NR5A2\|LHX4 | Arcuate Nucleus |
| C6-283 | NE_Mid-1_3_TBX3_PDGFD_GABRE_PGR | 211 | 63 | 29.86 | 1.24 | PGR\|CYSLTR2\|GABRE\|TBX3 | Arcuate Nucleus |
| C6-223 | NE_Mid-1_4_ADAMTSL1_FOXG1_SOX14_SHC4 | 47 | 14 | 29.79 | 0.24 | SKOR2\|CCN2\|ADCYAP1\|SHC4 | Low abundance |

C

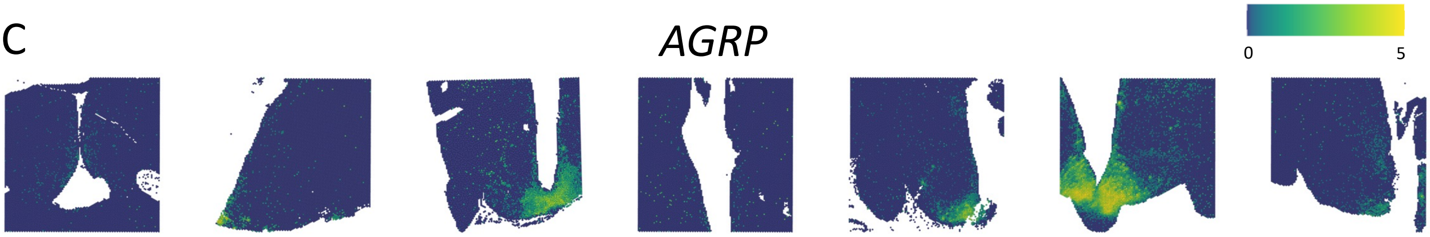


D

| Cluster | Cluster Name | Number of Cells | Number of AGRP+ cells | % AGRP+ | Average Expression | Cluster Marker Genes | Spatial Mapping |
|---|---|---|---|---|---|---|---|
| C6-260 | NE_Mid-1_3_OTP_SOX5_AGRP | 324 | 118 | 36.42 | 1.06 | AGRP\|AGTR1\|NPY\|OTP | Arcuate Nucleus |
| C6-212 | NE_Mid-1_2_2_GAL_GHRH_TENT5A | 184 | 43 | 23.37 | 0.61 | GHRH\|GAL\|ADGRF4\|ITGA4 | Arcuate Nucleus |
| C6-396 | NE_Mid-2_SIM1_STK32B_AVP_CARTPT_RORB | 152 | 28 | 18.42 | 1.38 | TH\|CARTPT\|AGRP\|TRH | Supraoptic Nucleus, Paraventricular Nucleus |
| C6-195 | NE_Mid-1_2_1_GAL_GPR101_ACVR1C | 599 | 101 | 16.86 | 0.26 | GAL\|MORC1\|MBNL3\|NTS | Arcuate Nucleus |
| C6-196 | NE_Mid-1_2_1_GAL_GPR101_GHRH | 132 | 20 | 15.15 | 0.14 | GAL\|GHRH\|TMEM155\|DLK1 | Arcuate nucleus |

E

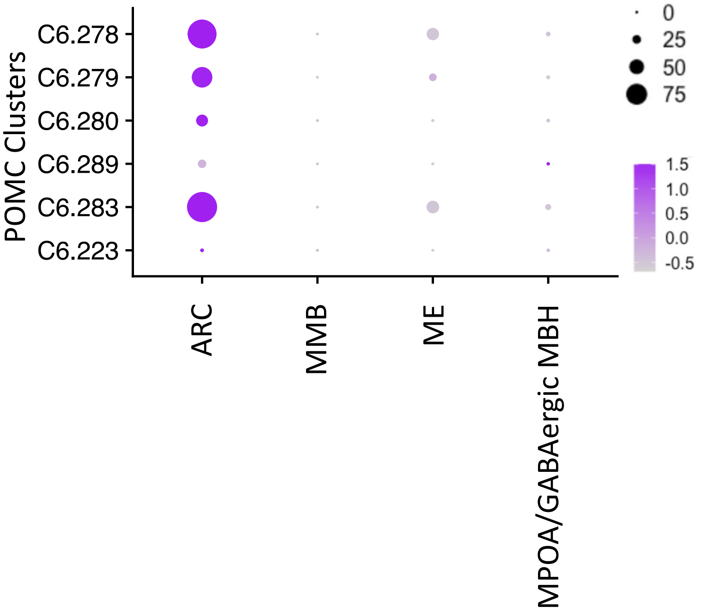


F

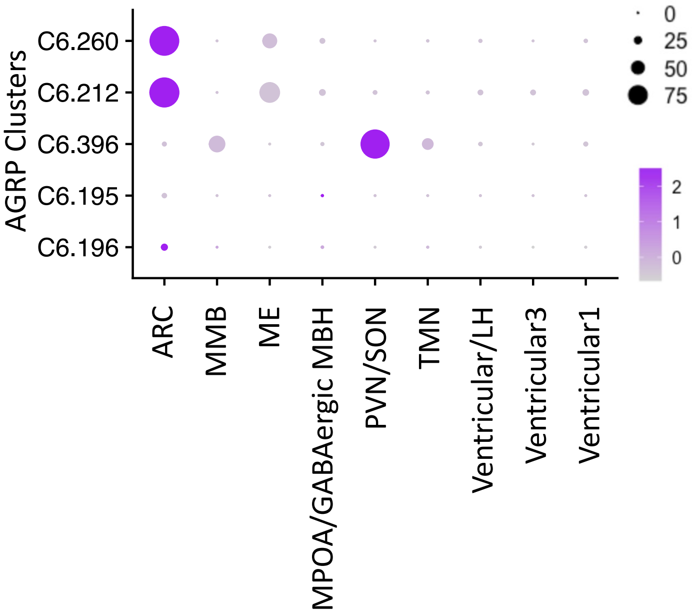

### Extended data 7: Characterisation of MC3R & MC4R populations

## *MC4R*

A

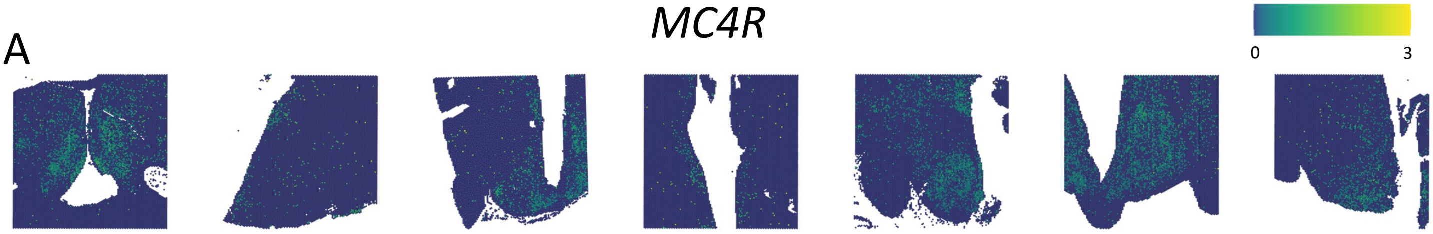


B

| Cluster | Cluster Name | Number of Cells | Number of MC4R+ cells | % MC4R+ | Average Expression | Cluster Marker Genes | Spatial Mapping |
|---|---|---|---|---|---|---|---|
| C6-43 | NE_Pre_1_MYO16_CXCL14_CAV1 | 110 | 54 | 49.09 | 0.49 | PROK2\|CAV1\|NTN1\|COL4A1 | Low Abundance |
| C6-253 | NE_Mid-1_4_PPP1R17_CALCRL_LEF1_PTGER3 | 21 | 7 | 33.33 | 0.19 | RASSF6\|ADGRG6\|HMCN1\|LEF1 | Low Abundance |
| C6-224 | NE_Mid-1_4_ADAMTSL1_FOXG1_CYP19A1_SKOR2 | 191 | 52 | 27.23 | 0.19 | SKOR2\|CYP19A1\|FAM9B\|QRFPR | Preoptic Area |
| C6-112 | NE_Pre_SLC44A5_SLC5A7_BMPR1B | 313 | 85 | 27.16 | 0.37 | SLC5A7\|COL6A5\|CHAT\|LHX8 | Lateral preoptic area |
| C6-407 | NE_Mid-2_SIM1_STK32B_NELL1_RASGEF1B_KCNH8 | 287 | 74 | 25.78 | 0.22 | OTP\|KCNH8\|LMCD1\|STK32B | Low Abundance |
| C6-181 | NE_Mid-1_2_FOXG1_HMX3_GLI3 | 215 | 50 | 23.26 | 0.20 | GLI3\|TPTE\|HMX3\|RORB | IMH |
| C6-295 | NE_Mid-1_3_LEF1_NRP2_PPP1R1C | 52 | 12 | 23.08 | 0.12 | LEF1\|PPP1R1C\|ANKRD30A\|TBX3 | Low Abundance |
| C6-276 | NE_Mid-1_3_1_WIF1_LEF1 | 32 | 7 | 21.88 | 0.24 | LEF1\|ETS1\|ADAMTS20\|KCNJ5 | Low Abundance |
| C6-88 | NE_Pre_SLC44A5_SFRP1_IL1RAPL2_SST | 191 | 41 | 21.47 | 0.23 | MOXD1\|LHX6\|IL1RAPL2\|NXPH2 | Low Abundance |
| C6-402 | NE_Mid-2_SIM1_STK32B_SLITRK6_NWD2_NPSR1 | 336 | 70 | 20.83 | 0.12 | NPSR1\|STK32B\|LYPD6\|SIM1 | Low Abundance |

## *MC3R*

C

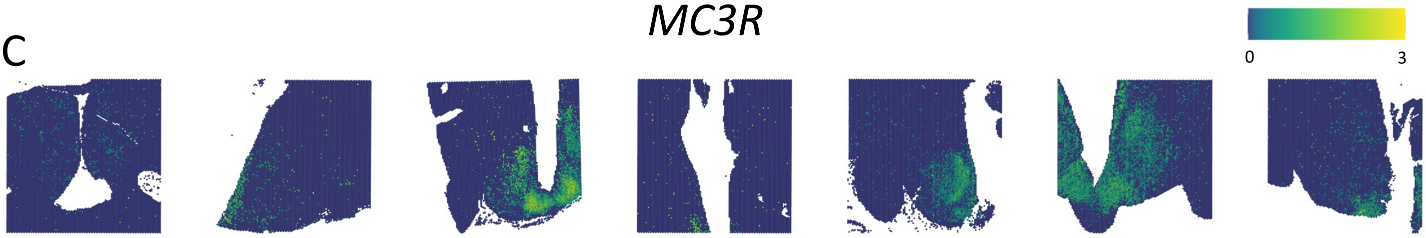


D

| Cluster | Cluster Name | Number of Cells | Number of MC3R+ cells | % MC3R+ | Average Expression | Cluster Marker Genes | Spatial Mapping |
|---|---|---|---|---|---|---|---|
| C6-289 | NE_Mid-1_3_TBX3_ESR1_CALCR | 143 | 33 | 23.08 | 0.06 | PGR\|CALCR\|NR5A2\|LHX4 | Arcuate Nucleus |
| C6-288 | NE_Mid-1_3_TBX3_ESR1_COL22A1 | 527 | 98 | 18.60 | 0.11 | KISS1\|SKOR2\|NR5A2\|PGR | Arcuate Nucleus |
| C6-196 | NE_Mid-1_2_1_GAL_GPR101_GHRH | 132 | 21 | 15.91 | 0.10 | GAL\|GHRH\|TMEM155\|DLK1 | Arcuate nucleus |
| C6-291 | NE_Mid-1_3_TBX3_ESR1_CCBE1_COL22A1 | 256 | 35 | 13.67 | 0.08 | KISS1\|SKOR2\|UGT2B7\|TAC3 | Arcuate Nucleus |
| C6-212 | NE_Mid-1_2_2_GAL_GHRH_TENT5A | 184 | 24 | 13.04 | 0.11 | GHRH\|GAL\|ADGRF4\|ITGA4 | Arcuate Nucleus |
| C6-283 | NE_Mid-1_3_TBX3_PDGFD_GABRE_PGR | 211 | 25 | 11.85 | 0.03 | PGR\|CYSLTR2\|GABRE\|TBX3 | Arcuate Nucleus |
| C6-243 | NE_Mid-1_4_ADAMTSL1_WDR49_FEZF1_CCBE1 | 989 | 111 | 11.22 | 0.09 | NR5A1\|COL15A1\|FEZF1\|CCBE1 | VMH |
| C6-280 | NE_Mid-1_3_TBX3_POMC_ANKRD30A | 384 | 40 | 10.42 | 0.04 | SOX3\|PGR\|POMC\|TBX3 | Arcuate Nucleus |
| C6-279 | NE_Mid-1_3_TBX3_POMC_CALCR | 351 | 27 | 7.69 | 0.03 | CALCR\|POMC\|WIF1\|PGR | Arcuate Nucleus |
| C6-125 | NE_Mid-1_1_SATB2_SOX6 | 483 | 29 | 6.00 | 0.10 | SATB2\|SLC6A3\|SATB2-AS1\|BRS3 | Arcuate Nucleus |

### Extended data 8: Incretin mapping

A

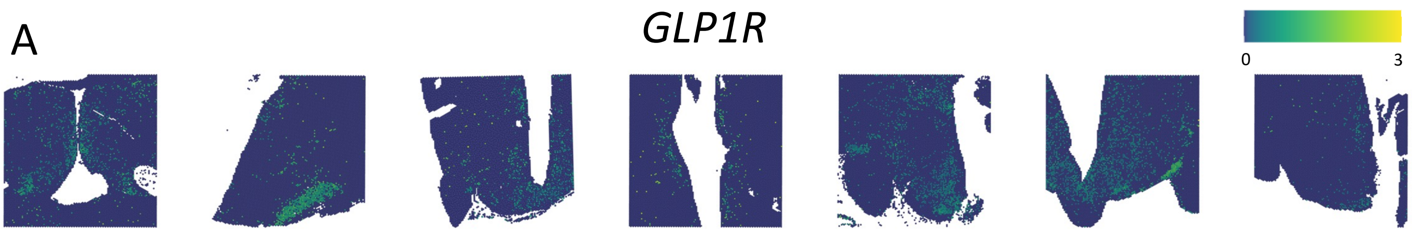


B

| Cluster | Cluster Name | Number of Cells | Number of GLP1R+ cells | % GLP1R+ | Average Expression | Cluster Marker Genes | Spatial Mapping |
|---|---|---|---|---|---|---|---|
| C6-414 | NE_Mid-2_SIM1_SCGN_EBF3_AVP | 344 | 268 | 77.91 | 1.11 | MEDAG\|FOXQ1\|SCGN\|TH | Supraoptic Nucleus, Paraventricular Nucleus |
| C6-209 | NE_Mid-1_2_2_GAL_CALCR_SST | 506 | 198 | 39.13 | 0.43 | CALCR\|SST\|SIX6\|DLK1 | MBH (outside arc) |
| C6-84 | NE_Pre_SLC44A5_SFRP1_SULF1_CTXND1 | 853 | 281 | 32.94 | 0.85 | CER1\|CTXND1\|TAC3\|NXPH2 | Low Abundance |
| C6-104 | NE_Pre_SLC44A5_BCL11A_INPP4B_ETV1 | 126 | 33 | 26.19 | 0.27 | GBX1\|LHX8\|ELFN1\|MEGF10 | Low Abundance |
| C6-413 | NE_Mid-2_SIM1_SCGN_EBF3_HS3ST2 | 342 | 88 | 25.73 | 0.35 | TH\|VWA3B\|HS3ST2\|PPP1R17 | Supraoptic Nucleus, Paraventricular Nucleus |
| C6-396 | NE_Mid-2_SIM1_STK32B_AVP_CARTPT_RORB | 152 | 36 | 23.68 | 0.43 | TH\|CARTPT\|AGRP\|TRH | Supraoptic Nucleus, Paraventricular Nucleus |
| C6-285 | NE_Mid-1_3_TBX3_PDGFD_PIK3C2G_WIF1 | 502 | 117 | 23.31 | 0.23 | PIK3C2G\|RERGL\|WIF1\|TBX3 | Median Eminence |
| C6-278 | NE_Mid-1_3_TBX3_POMC_LEPR | 414 | 91 | 21.98 | 0.14 | POMC\|WIF1\|GABRE\|TBX3 | Arcuate Nucleus |
| C6-374 | NE_Mid-2_ZIC2_THSD7B_TBR1_EBF2 | 82 | 16 | 19.51 | 0.37 | EBF2\|EBF3\|TNC\|EOMES | Low Abundance |
| C6-198 | NE_Mid-1_2_1_TCF7L1_ROR1 | 464 | 86 | 18.53 | 0.28 | TMEM176B\|FOXD2\|PENK\|GAL | Periventricular hypothalamus |

C

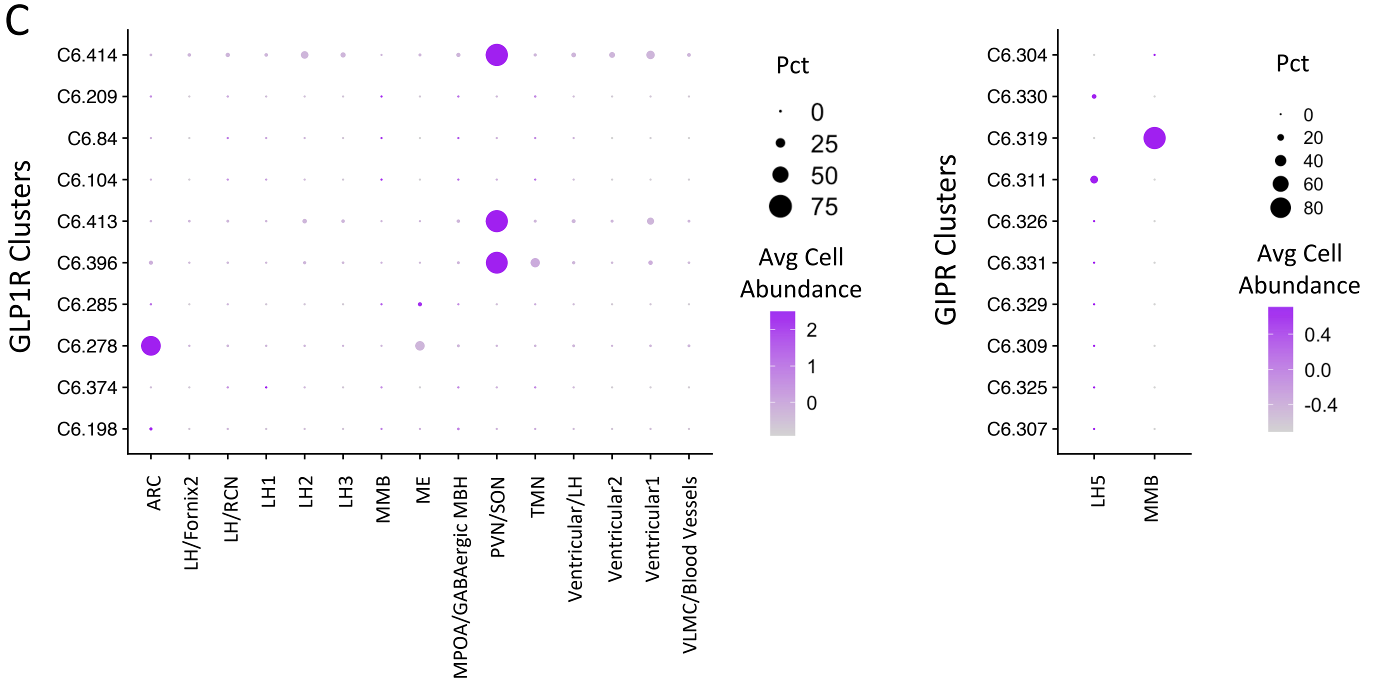


D

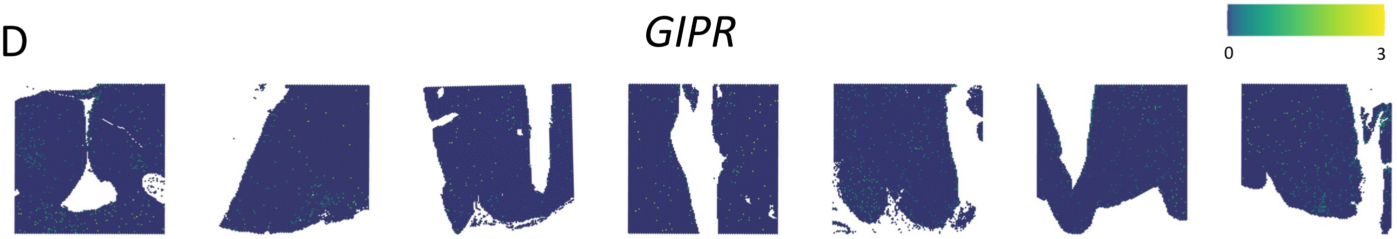


E

| Cluster | Cluster Name | Number of Cells | Number of GIPR+ cells | % GIPR+ | Average Expression | Cluster Marker Genes | Spatial Mapping |
|---|---|---|---|---|---|---|---|
| C6-304 | NE_LMX1A_PDZRN4_EBF2_CRNDE_PLEKHH2 | 67 | 42 | 62.69 | 0.72 | EBF2\|PLEKHH2\|TTC6\|LMX1A | Low Abundance |
| C6-330 | NE_LMX1A_NTNG2_COBLL1_NEAT1 | 639 | 342 | 53.52 | 0.40 | LMX1A\|CDH23\|COBLL1\|AP001636.3 | Lateral hypothalamus 5 |
| C6-319 | NE_LMX1A_CRNDE_PTPRK_EBF2 | 378 | 201 | 53.17 | 0.48 | CRNDE\|LMX1A\|CARTPT\|TTC6 | MMB |
| C6-311 | NE_LMX1A_PDZRN4_ZEB2_EBF2_TTC6 | 493 | 258 | 52.33 | 0.41 | AP005242.1\|C1QL1\|EBF2\|LMX1A | Lateral hypothalamus 5 |
| C6-326 | NE_LMX1A_NTNG2_NTS_CDH23_ANOS1 | 162 | 69 | 42.59 | 0.30 | NTS\|EBF2\|FAM83B\|CDH23 | Low Abundance |
| C6-331 | NE_LMX1A_NTNG2_COBLL1_FBN2 | 80 | 32 | 40 | 0.36 | NTS\|LMX1A\|CDH23\|LMX1B | Low Abundance |
| C6-329 | NE_LMX1A_NTNG2_EBF2 | 344 | 134 | 38.95 | 0.35 | EBF2\|LMX1A\|EBF3\|BMP5 | Low Abundance |
| C6-309 | NE_LMX1A_PDZRN4_ZEB2_ITGA8_VIP | 611 | 234 | 38.298 | 0.38 | VIP\|LMX1A\|TTC6\|AP005242.1 | Low Abundance |
| C6-325 | NE_LMX1A_NTNG2_NTS_CDH23_EBF2 | 330 | 119 | 36.06 | 0.31 | EBF2\|NTS\|LMX1A\|FSHR | Low Abundance |
| C6-307 | NE_LMX1A_PDZRN4_EBF2_CRNDE_POSTN | 131 | 42 | 32.06 | 0.31 | POSTN\|SLC26A7\|EBF2\|LMX1A | Low Abundance |

### Extended data figure 9

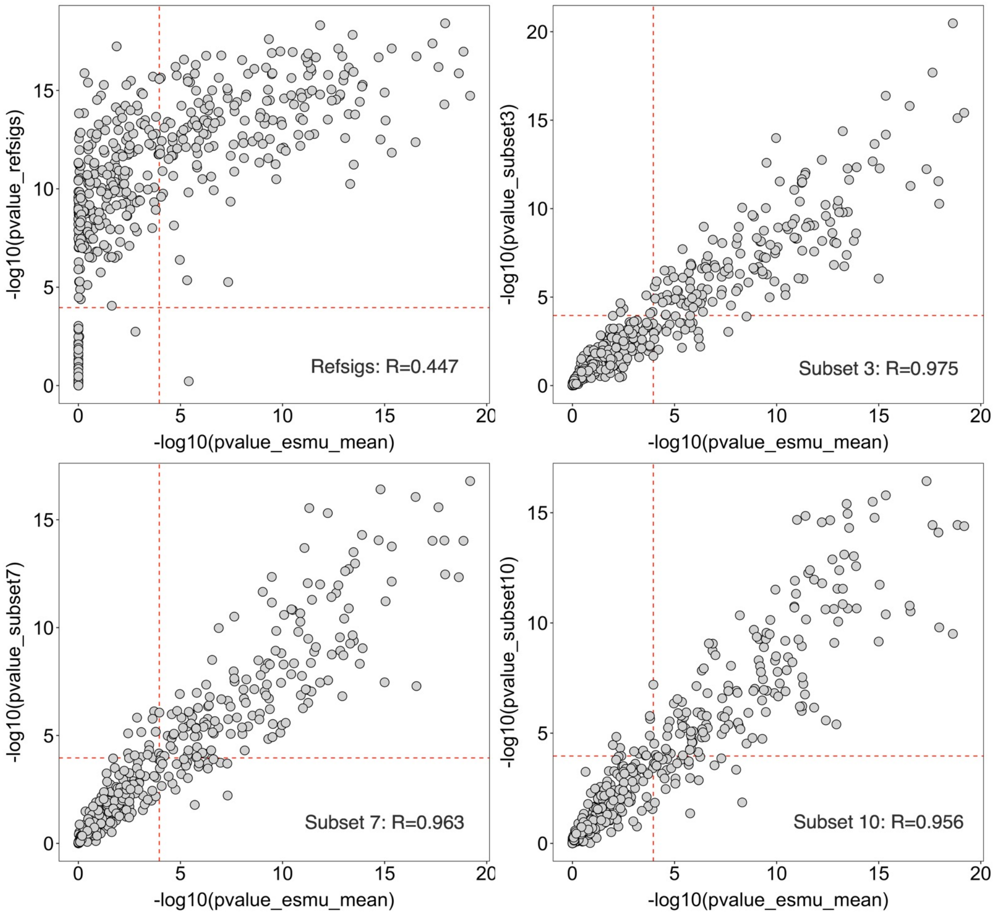
